# Impacts of disease surveillance frequency on understanding and controlling Rift Valley fever virus

**DOI:** 10.1101/2025.08.11.669600

**Authors:** Warren S. D. Tennant, Jake Carson, Glen Guyver-Fletcher, Raphaëlle Métras, Michael J. Tildesley

## Abstract

Effective livestock disease surveillance faces numerous socio-economic and disease-specific challenges, particularly within resource-limited settings. Rift Valley fever (RVF), a zoonotic vector-borne disease which is of growing threat to human and veterinary health, exemplifies these challenges, consequently leading to irregular or infrequent surveillance. While mathematical models frequently utilise these surveillance data to understand disease dynamics and assess the effectiveness of disease control measures, the impact of surveillance frequency on these model-based assessments is not clear. To address this, we used a livestock RVF virus infection model, which incorporates spatial and age structures, livestock movement, and environmental factors. We then generated synthetic cross-sectional seroprevalence surveys with varying frequencies from outbreak scenarios inferred from empirical serological data from the Comoros archipelago. By refitting the model to these synthetic data, we found that lower surveillance frequency was associated with increased uncertainties in understanding disease biology, inferring past outbreak timing, predicting future outbreak size, and recommending optimal control measures. In particular, once serological surveillance was less frequent than annually, the ability to distinguish spatio-temporal patterns of disease transmission and forecast trends diminished. These findings emphasise the need for adequately frequent surveillance data to ensure robust model-based analyses which inform effective preparedness and control strategies for livestock diseases within resource-limited settings.

**Author summary:** Disease surveillance underpins mathematical models that are used to understand the emergence, spread and persistence of disease, and assess and compare the impacts of disease control measures. However, regular surveillance is a particular challenge for livestock diseases in resource-limited settings, such as those affected by Rift Valley fever. Here, our work provides an illustrative example of how surveillance frequency impacts our ability to understand disease outbreaks and to decide which control measures are most effective at combatting disease, using outbreak scenarios for Rift Valley fever in the Comoros archipelago. By simulating from a mathematical model informed by different surveillance frequencies, we found that less frequent surveillance was associated with greater uncertainty in model predictions, and demonstrated that this may distort epidemiological understanding and future control recommendations of a disease. Our work has crucial implications for animal health decision-making, highlighting the importance of consistent and frequent surveillance to improve disease management and prioritise disease control measures.

## Introduction

Designing effective surveillance systems for livestock diseases presents numerous challenges [1–5]. Socio-economic factors, such as resource limitations [1], inadequate knowledge of disease [2, 4], loss of income [6, 7] or social stigma associated with disease notification [8], lack of trust in government [9, 10], and absence of standardisation in data systems [5], hinder or discourage disease reporting. Asymptomatic or subclinical infections, wildlife reservoirs of infection [3, 11], sensitivity and specificity of diagnostic tests [4, 12], and spatio-temporal variations in disease transmission and livestock demography [13–15], can further obscure the true extent of disease circulation. Rift Valley fever (RVF) — a zoonotic viral haemorrhagic fever primarily affecting livestock in lower-middle-income countries — exemplifies these difficulties. RVF outbreak detection relies on either observing rare, extreme events like mass abortions in ewes [16, 17], or through serological evidence of exposure to the virus [18–20]. Moreover, surveillance of RVF is often dependent on time-limited funding or national programs [19] that are susceptible to competing disease priorities between regions [21–23]. These limiting factors compound in resource-limited settings, and consequently may result in infrequent or irregular disease monitoring across space and time.

Livestock disease surveillance data is frequently integrated within mathematical models to generate insights into disease dynamics [15, 24–27] and evaluate the impact of control measures [28–32]. However, the degree to which these model-derived insights are contingent on the quality and consistency of the underlying surveillance data is not always clear. While studies have shown that the increased surveillance data as an outbreak progresses improves confidence in disease forecasts and control recommendations [32–35], there has yet to be a critical quantitative evaluation of how surveillance frequency impacts our understanding of RVF and control policy decisions.

To illustrate the impact of surveillance frequency on model-based inference for RVF, we first modified an existing livestock Rift Valley fever virus (RVFV) infection model [15]—which incorporates age and spatial structure, movement of livestock and environmental factors influencing disease transmission rates—to include natural randomness in disease transmission and allow for stochastic extinction of the disease. This model was fitted in a Bayesian framework to age-stratified serological data from July 2008 to June 2015 collected across the Comoros archipelago (Fig 1A): a network of four islands (Grande Comore, Mohéli, Anjouan and Mayotte) in the south-western Indian Ocean. This fitted model was used to infer realistic outbreak scenarios (Fig 1B), from which synthetic serological surveys were generated, each with a different surveillance frequency (Fig 1C). The model was then fitted back to each of these datasets (Fig 1D) and forecasts were performed for an additional three years under a range of disease control strategies. This procedure allowed for a comprehensive demonstration of how surveillance frequency impacts the ability to understand past outbreaks, forecast future trends, and prioritise disease control measures.

**Fig 1.**
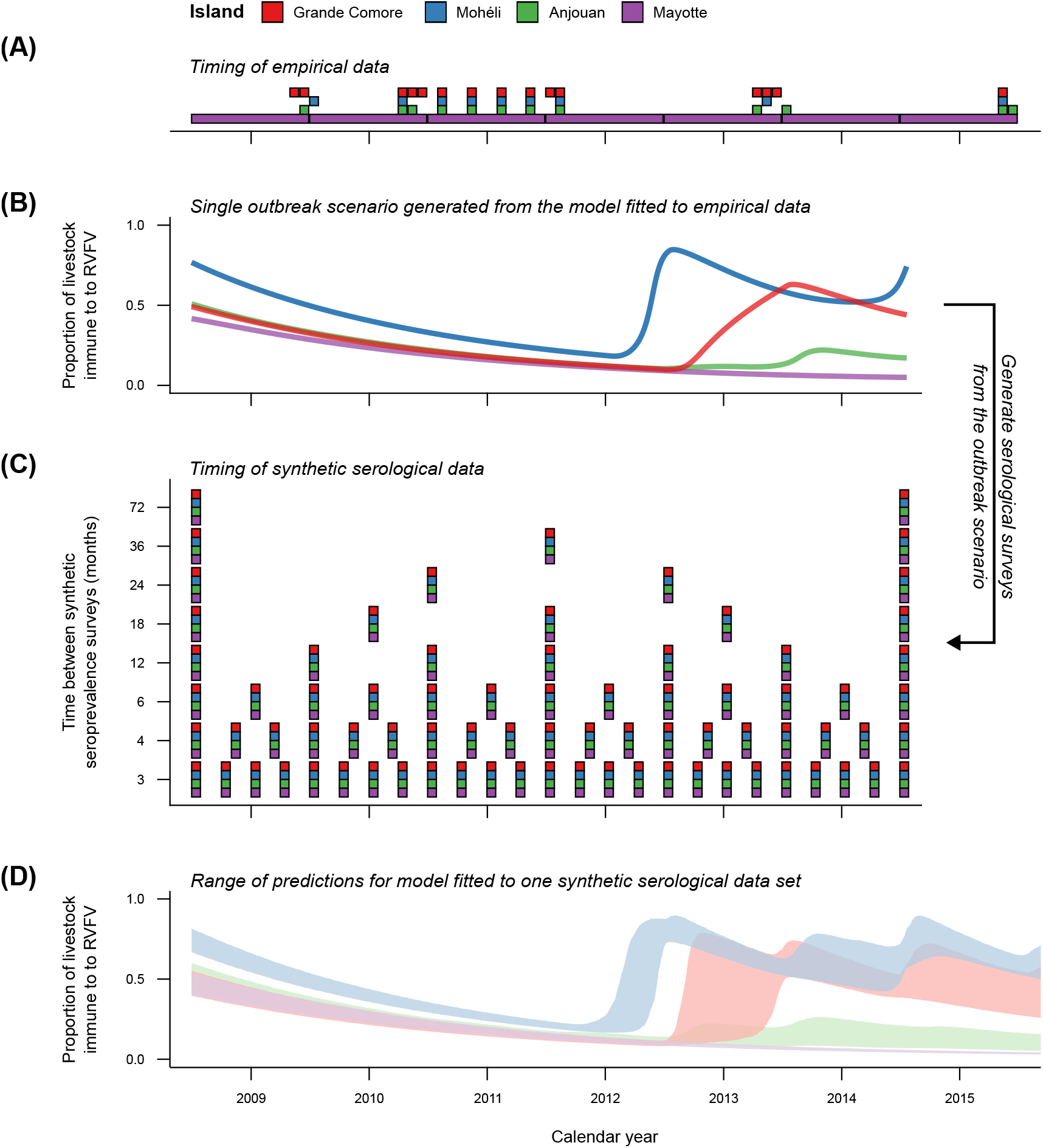
Schematic for the assessment of surveillance frequency effects on model-based decision making. **(A)** The timing and duration of empirical seroprevalence surveys which were conducted between 2008 and 2015 on each island in the Comoros archipelago—Grande Comore (red), Mohéli (blue), Anjouan (green) and Mayotte (purple). **(B)** Realistic outbreak scenarios, as illustrated by weekly model-predicted seroprevalence on each island, were inferred by fitting the mathematical model to this empirical data in a Bayesian framework. **(C)** For each of five outbreak scenarios, synthetic serological data were generated with eight different surveillance frequencies as shown, yielding a total of 40 data sets. **(D)** The model was then fitted to each of these data sets and simulated forward in time under a range of control scenarios to quantify the impact of surveillance frequency on forecasts and model-based decision making. Shown is the range of historical and forecasted predictions of weekly seroprevalence on each island with annual seroprevalence surveys in the absence of controls.

## Results

Five RVF outbreak scenarios were extracted from the livestock infection model, which was fitted in a Bayesian framework to empirical data from the Comoros archipelago. Each scenario exhibited outbreaks of RVF in Grande Comore, Mohéli and Anjouan as shown by rises in model-predicted seroprevalence, representing the proportion of livestock with prior RVFV infection, between July 2008 and June 2015 (S1 Fig). Synthetic one month-long age-stratified serological surveys were created every 3 months, 4 months, 6 months, 1 year, 18 months, 2 years, 3 years and 6 years for each outbreak scenario. The main results below present one of the five outbreak scenarios.

### Biological inference

Island-specific disease introduction and transmission rates, the initial proportion of the livestock population immune to infection, and levels of heterogeneity in livestock viral exposure, were estimated by fitting the mathematical model to the synthetic data with different surveillance frequencies. S1 Data presents the median and 95% credible interval (CrI) of these estimates for all surveillance frequencies and outbreak scenarios.

We first found that the estimates recovered the values used to generate each synthetic data set across all surveillance frequencies (S2–S3 Fig). We also observed an increase to the uncertainty in these estimates as surveillance frequency decreased. For example, the expected reproduction number of the disease — the mean number of secondary infections from a single primary infection in a fully susceptible population — was 1.68 on Grande Comore (Table 1). This value was within the 95% credible interval of reproduction number estimates for seven of the eight analysed surveillance frequencies, with the intervals widening as surveillance frequency decreased. Specifically, the median reproduction number estimate was 1.59 (95% CrI = [1.46, 1.73]) when surveys were conducted every 3 months, 1.49 (95% CrI = [1.32, 1.72]) for surveys every six months, 1.85 (95% CrI = [1.35, 2.36]) for surveys every year, 1.39 (95% CrI = [1.26, 1.82]) for surveys every two years, and 1.84 (95% CrI = [1.39, 2.5]) for surveys every six years in Grande Comore.

**Table 1.** Reproduction number estimates at different surveillance frequencies. Synthetic serological surveys were generated for every 3 months, 4 months, 6 months, 1 year, 18 months, 2 years, 3 years and 6 years using outbreak scenarios inferred from empirical data. The table shows the median and 95% credible interval of reproduction number estimates—the expected number of secondary infections from a single infection in an otherwise susceptible population—on each island for five surveillance frequencies and a single outbreak scenario. The uncertainty in these estimates increased as surveillance frequency decreased on each island. Summary metrics were calculated using 2500 samples from the posterior distribution of each surveillance frequency.

| Island | Time period between synthetic seroprevalence surveys (months) | Median reproduction number [95% CrI] |
| --- | --- | --- |
| Grande Comore | Outbreak scenario | 1.68 |
|  | 3 | 1.59 [1.46, 1.73] |
|  | 6 | 1.49 [1.32, 1.72] |
|  | 12 | 1.85 [1.35, 2.36] |
|  | 24 | 1.39 [1.26, 1.82] |
|  | 72 | 1.84 [1.39, 2.50] |
| Mohéli | Outbreak scenario | 2.69 |
|  | 3 | 2.67 [2.45, 2.99] |
|  | 6 | 2.95 [2.59, 3.42] |
|  | 12 | 2.83 [2.42, 3.59] |
|  | 24 | 2.52 [1.94, 3.53] |
|  | 72 | 2.51 [1.53, 3.85] |
| Anjouan | Outbreak scenario | 1.16 |
|  | 3 | 1.16 [1.10, 1.21] |
|  | 6 | 1.17 [1.09, 1.25] |
|  | 12 | 1.14 [1.00, 1.23] |
|  | 24 | 1.18 [1.07, 1.39] |
|  | 72 | 1.31 [1.10, 1.67] |
| Mayotte | Outbreak scenario | 0.51 |
|  | 3 | 0.13 [0.00, 0.77] |
|  | 6 | 0.11 [0.00, 0.83] |
|  | 12 | 0.12 [0.00, 0.84] |
|  | 24 | 0.13 [0.01, 0.94] |
|  | 72 | 0.14 [0.00, 1.10] |

The extent of the uncertainty in epidemiological estimates depended on both the parameter and the island for which they were estimated. For instance, the expected annual number of disease introductions (infections not attributed to locally infected livestock) was 1.04 in Mohéli and 19.73 in Mayotte. On Mohéli, the 95% credible interval for this parameter widened as surveillance frequency decreased beyond one survey every year: [0.04, 1.56], [0.03, 1.24], [0.03, 1.57], [0.05, 3.89] and [0.40, 25.50] for surveys every three months, six months, year, two years and six years. In contrast, Mayotte’s intervals showed minimal variation: [0.37, 25.16], [0.41, 27.26], [0.30, 27.25], [0.62, 28.47] and [0.35, 27.78] for the respective surveillance frequencies. In Mayotte, these intervals closely reflected the range of uncertainty present before any data was considered (the prior distribution), suggesting insufficient data to improve the estimate. This may be attributed to the lack of outbreaks in Mayotte in the outbreak scenario.

### Timing of historical outbreaks

These findings translated across to predictions of livestock population immunity levels over time. That is, the outbreak scenario itself was within the variability of modelled seroprevalence, and the uncertainty in seroprevalence increased as surveillance frequency decreased. Fig 2 shows the model-inferred RVFV-specific immunoglobulin-G (IgG) seroprevalence over time under each surveillance frequency.

**Fig 2.**
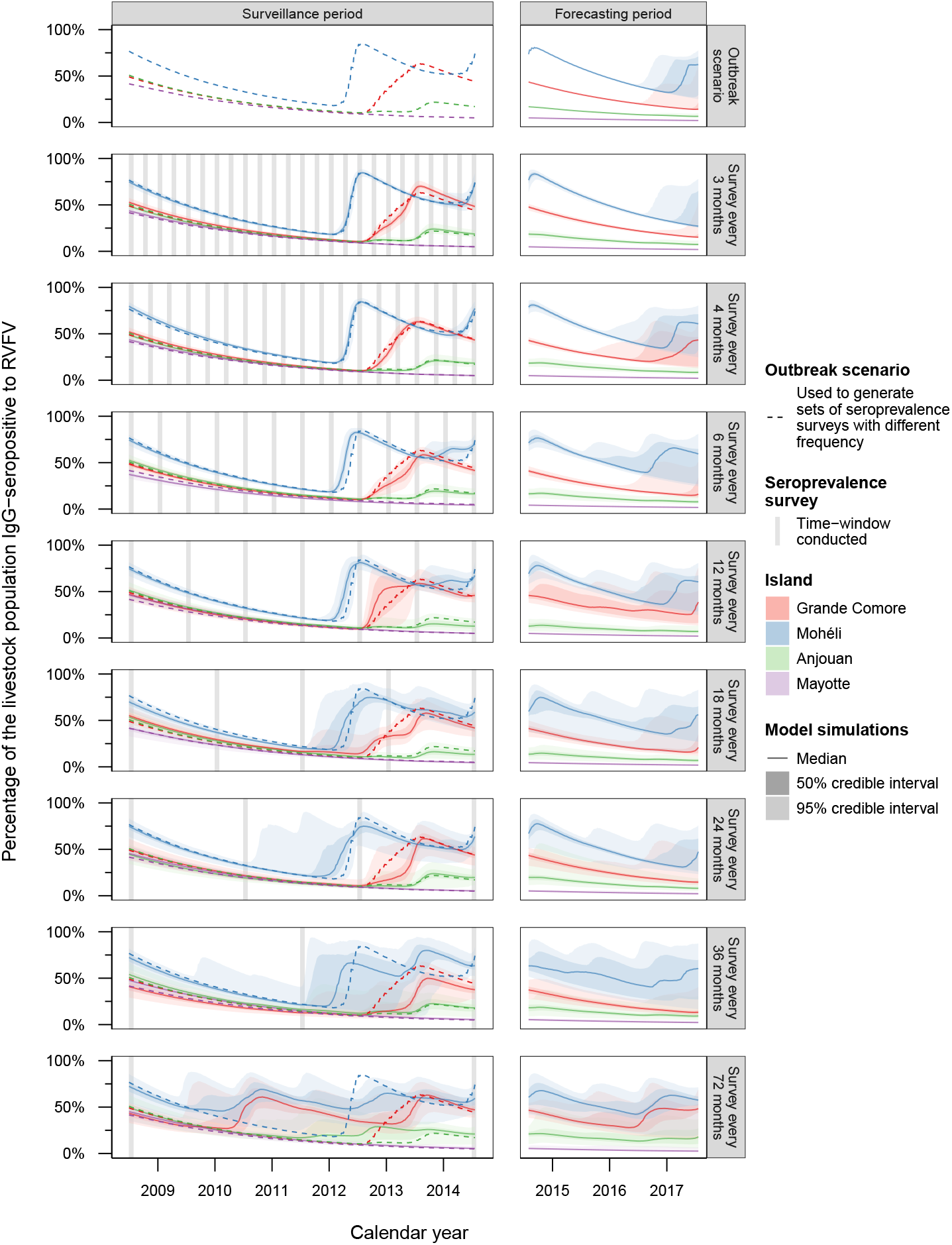
Historical and forecasted IgG-seroprevalence to RVFV. An outbreak scenario (dashed line) was generated by fitting a mathematical model to empirical age-stratified seroprevalence surveys conducted across the archipelago from July 2008 to June 2015. From this, synthetic serological surveys were generated with different intervals (vertical grey bars): from once every 3 months to once every 6 years. The model was fitted independently to each of these data. Shown is the median (solid line), 50% credible interval and 95% credible interval (shaded ribbons) of the percentage of livestock with past exposure to RVFV on each island in the Comoros arhcipelago—Grande Comore (red), Mohéli (blue), Anjouan (green) and Mayotte (purple). As surveillance frequency decreased, the uncertainty in predicted IgG-seroprevalence between surveys increased across Grande Comore, Mohéli and Anjouan, which was carried through to model-based forecasts after surveillance had ended. The medians and credible intervals were based on 2500 posterior samples and corresponding forward-simulations for each surveillance frequency.

For all surveillance frequencies, the expected population-level seroprevalence closely matched the modelled seroprevalence at the times surveys were conducted. For example, the expected seroprevalence in July 2008 was 0.49, 0.76, 0.51, 0.42 in Grande Comore, Mohéli, Anjouan and Mayotte, respectively. When surveys were conducted every 3 months, the median estimates were 0.53 in Grande Comore (95% CrI = [0.48, 0.58]), 0.75 in Mohéli (95% CrI = [0.70, 0.79]), 0.49 in Anjouan (95% CrI = [0.44, 0.54]), 0.44 in Mayotte (95% CrI = [0.40, 0.47]). If surveys were only conducted every six years, these estimates were 0.43 in Grande Comore (95% CrI = [0.32, 0.54]), 0.72 in Mohéli (95% CrI = [0.55, 0.85]), 0.49 in Anjouan (95% CrI = [0.35, 0.63]), and 0.45 in Mayotte (95% CrI = [0.35, 0.55]).

Decreasing surveillance frequency further led to increased variability in inferred seroprevalence between serological surveys (Figure 2). This corresponded to increased uncertainty over when and how many outbreaks had occurred during the surveillance period. For instance, on Mohéli, when surveillance was conducted quarterly to annually, two major outbreaks—defined as greater than 10% of livestock infected in a single epidemiological year—were correctly identified in 2011/12 and 2013/14. Additionally, the within-year timings of these outbreaks were more accurately inferred as surveillance frequency increased. However, when the interval between surveys increased beyond 18 months, the inferred timing of outbreaks was found to be less precise, with the model suggesting a possible additional outbreak in the 2012/13. Reducing surveillance to only once every six years, the model indicates that outbreaks may have occurred in any epidemiological year, with a median of 4 major outbreaks (95% prediction interval (PrI) = [2, 6]) across the surveillance period. Grande Comore and Anjouan exhibited similar trends to Mohéli, whilst Mayotte, which demonstrated no major outbreaks in the outbreak scenario, showed robust seroprevalence decay irrespective of surveillance frequency. These findings were consistent across all analysed outbreak scenarios (S4–S7 Fig).

### Forecasting future trends

Uncertainty in RVFV outbreak timing during the surveillance period extended into the 3-year forecasts (Figure 2). When surveillance was frequent, forecasts predicted a single outbreak event in Grande Comore and Mohéli, occurring two to three years after the surveillance period ended. As survey frequency decreased, the timing and the magnitude of outbreaks exhibited increased variability. Fig 3 illustrates the sensitivity of surveillance frequency on the total number of RVFV livestock infections on each island for each forecasting year.

**Fig 3.**
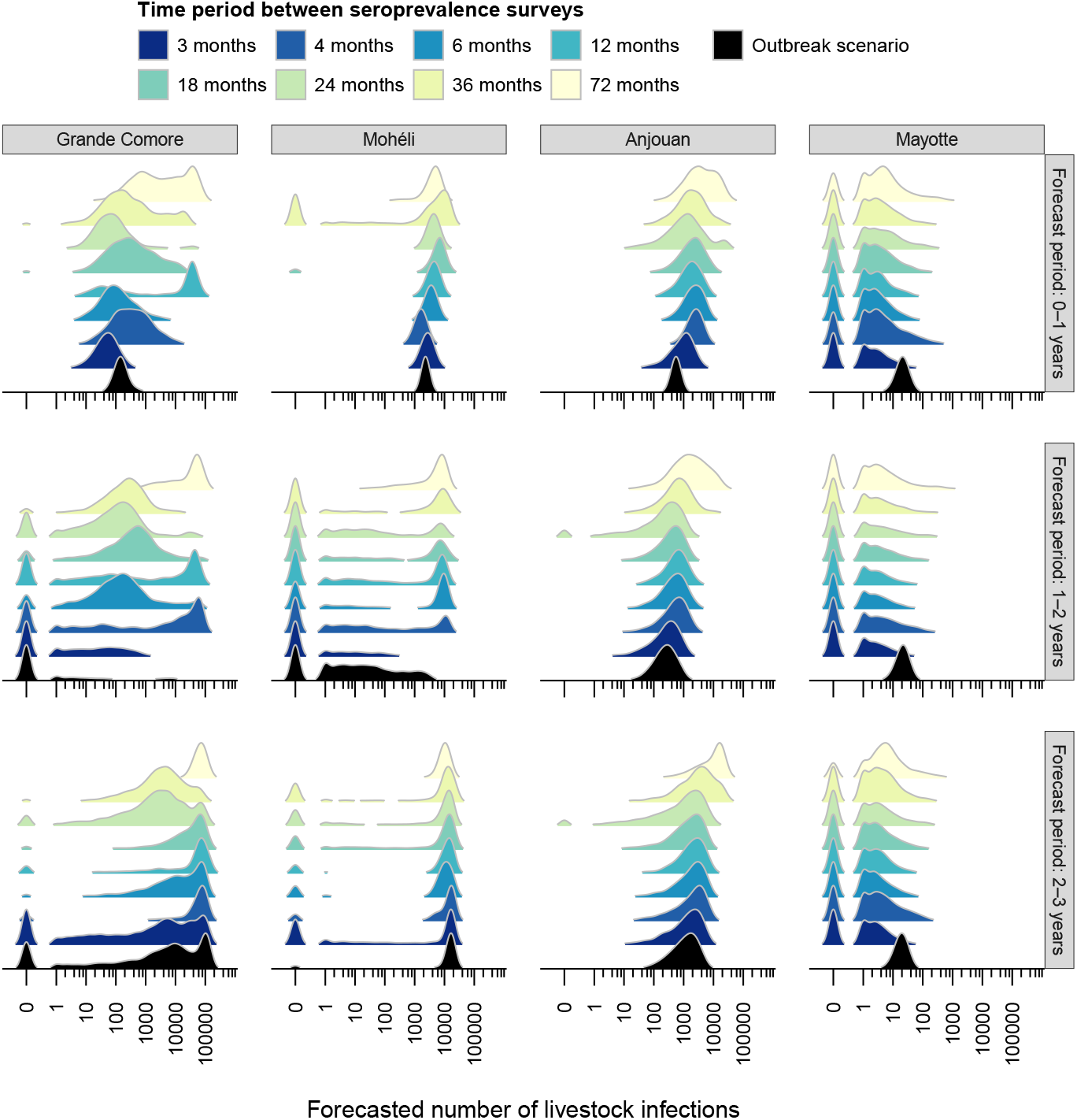
Surveillance frequency influence on outbreak size forecasts. The model was simulated forward in time for three years without disease control measures. Shown is the predicted number of livestock infections for each year following the end of disease surveillance, which were conducted every 3 months (dark blue) to six years (light yellow). The estimated distribution of infections from the outbreak scenario are shown in black. The forecasted number of infections in Anjouan and Mayotte appeared to be consistently predicted across different surveillance frequencies. However, Grande Comore and Mohéli exhibited variation in predicted outbreak size across surveillance frequency. Estimates shown were based on 2500 simulations for each surveillance frequency from a single outbreak scenario.

The influence of surveillance frequency on the robustness of forecasted livestock infections varied across the islands. Anjouan and Mayotte displayed relatively stable forecasts, with the magnitude of predicted outbreaks showing limited variation despite changes in surveillance frequency. Specifically, in Anjouan’s first forecast year, median livestock infections were: 560 (95% PrI = [367, 839]) with quarterly surveillance, 960 (95% PrI = [39, 3707]) with annual surveillance, 1726 (95% PrI = [162, 6939]) with biennial surveillance, and 3628 (95% PrI = [142, 26316]) with surveillance every six years. In contrast, Grande Comore and Mohéli showed marked variability in predicted outbreak size, with forecasts spanning from no outbreaks to major outbreaks, demonstrating a strong dependence on surveillance frequency.

In Grande Comore, the predicted probability of an outbreak—defined as when greater than 1% of livestock are infected—in the first forecasting year demonstrated a high sensitivity to surveillance frequency. Specifically, the probabilities were: 0.00 (quarterly), 0.04 (biannually), 0.57 (annually), 0.06 (biennially), 0.20 (triennially), and 0.54 (six-yearly) with an expected probability of 0.00 under the outbreak scenario. This initial variability propagated through to subsequent forecasting years. This was because the occurrence and magnitude of outbreaks in later years were dependent on the level of immunity conferred by outbreaks in the previous years, with larger initial outbreaks leading to reduced probabilities of later outbreaks. In the final forecast year, outbreak probabilities in Grande Comore ranged from 0.47 (quarterly) to 0.98 (six-yearly) with an expected probability of 0.56 under the outbreak scenario.

These characteristics were observed in the other four outbreak scenarios as well (S8 Fig). However, Anjouan was more adversely affected by surveillance frequency in these alternate outbreak scenarios compared with the one presented here in the main text. This may be due to stochastic variations in outbreak dynamics across scenarios.

### Control measure decision-making

Three-year forecasts demonstrated that the predicted epidemiological impact of disease control measure— livestock movement restrictions, reduction to disease transmission (e.g. vector control) and livestock vaccination— was also influenced by the frequency of surveillance in the past. Specifically, the predicted number of livestock infections of each disease control measure showed increased uncertainty as surveillance frequency decreased (S9 Fig). S1 Data presents the median and 95% credible interval of the predicted number of infections for all control measures, surveillance frequencies and outbreak scenarios.

To highlight how surveillance frequency may influence the control decision making process, we ranked the controls according to the predicted number of livestock infections. Fig 4 shows these rankings and the percentage of times each control measure achieved the highest rank. We found that the ranking of control strategies on all islands was sensitive to the frequency of surveys (Fig 4A). We observed that within each control type, stronger measures consistently ranked higher than weaker ones. However, the control rankings of all surveillance frequencies differed from that of the outbreak scenario. This is attributable to the higher uncertainty inherent in surveillance data, compared to the outbreak scenario’s perfect information, where disease transmission rates and the immunological status of all animals are fully known.

**Fig 4.**
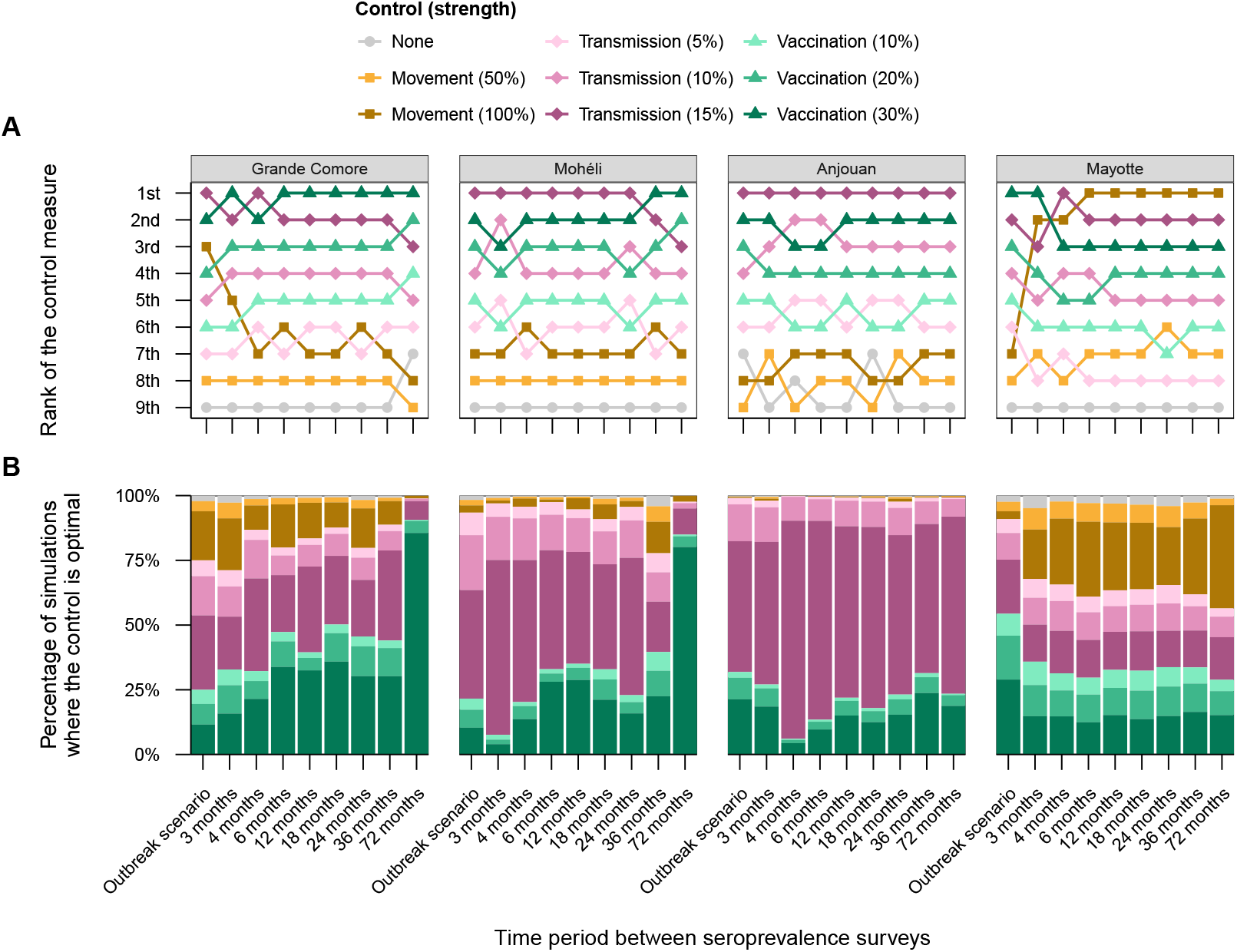
Influence of surveillance frequency on confidence in disease control measure forecasts. For each surveillance frequency and outbreak scenario, the number of livestock infections over a 3-year period was forecasted without (grey) and with different control measures: movement (yellow squares), reduction to transmission (pink diamonds) and vaccination (green triangles). **(A)** The control strategies were ranked based upon the total infections predicted within each island (lower predictions receiving higher ranks), where the ability to identify true ranking (as in the outbreak scenario) changed with different frequency of disease surveillance. **(B)** This effect was reflected in the proportion of simulations where each control strategy was the best (ranked first), with decreased confidence as surveillance frequency decreased. Model forecasts were based on 2500 simulations for each control measure and surveillance frequency. These are not policy control recommendations for the Comoros archipelago, but illustrate that any recommendations are sensitive to the surveillance of frequency underpinning them. Refer to S9 Fig for the predicted number of livestock infections for each scenario.

The frequency of surveys during the surveillance period also influenced the confidence level associated with the forecasted optimal control measure on each island (Fig 4B). For example, with quarterly surveys, vaccination of livestock was the most effective choice in only 15% and 4% of simulations in Grande Comore and Mohéli respectively. In contrast, when surveys were only conducted every six years, 85% and 80% of simulations showed that vaccination was the best control strategy, respectively. This shift may stem from the increased uncertainty associated with reproduction number estimates on these islands for low surveillance frequencies (see Table 1). In contrast, Anjouan and Mayotte exhibited greater stability in the confidence levels associated with the optimal control across surveillance frequency. In Anjouan, this stability may be attributed to lower reproduction number estimates which were reduced below the critical outbreak threshold (reproduction number equal to one) under reductions to disease transmission. As a result, the proportion of simulations where transmission reduction was the best ranged from 55% (quarterly) to 84% (every four months). In Mayotte, where no outbreaks were predicted regardless of surveillance frequency, the impact of different control strategies was comparable. Hence, the distribution of optimal strategies was robust to different surveillance frequencies.

The sensitivity of control strategy outcomes to surveillance frequency was observed in the other four outbreak scenarios (S10 Fig and S11 Fig). An exception to this was the control measure ranking for Mohéli in the final outbreak scenario, which demonstrated complete invariance to surveillance frequency. This may be attributed low seasonal variability in disease transmission rates in this specific outbreak scenario (S3 Fig). The reduced variability consequently may have improved the stability and predictability of disease outbreaks in Mohéli across all surveillance frequencies (S7 Fig) and control measures.

## Discussion

Robust disease surveillance is fundamental to understanding and controlling disease. However, irregular or untargeted surveillance can hinder our ability to detect outbreaks, comprehend disease dynamics, and forecast future trends. Livestock diseases pose unique challenges due to diverse livestock management practices, reduced knowledge of circulating diseases and lack of standardised reporting systems. Furthermore, zoonotic diseases, such as Rift Valley fever, often present subclinically and may thus rely on detection through serological monitoring or freak events such as abortion storms. These challenges are particularly pronounced in resource-limited settings where regular surveillance may be infeasible. While model-based approaches can elucidate the factors responsible for disease spread, as well as predict the impact of control measures, the influence of surveillance frequency itself on our understanding and recommendations of livestock disease is not fully realised. To address this, we employed a model-based approach to assess how the frequency of cross-sectional serological testing impacts disease comprehension, outbreak detection, and disease control recommendations, using Rift Valley fever in the Comoros archipelago as a case study.

We consistently observed that decreased frequency of surveillance increased the uncertainty associated with (i) understanding the biology of the disease, (ii) inferring the timing of past outbreaks, (iii) predicting the size of future outbreaks and (iv) recommending optimal disease control measures.

We first found that estimates of epidemiological parameters became more uncertain as surveillance frequency decreased. A primary purpose of parameter estimation is to quantify the influence of disease transmission mechanisms on disease emergence, spread, and persistence. This included estimates for the basic reproduction number, a key measure of disease transmissibility and an indicator of the population immunity threshold required to prevent outbreaks. Greater uncertainty in these estimates will thus impact the precision of estimates for the level of immunity needed to avert an outbreak through control measures such as vaccination, one of the main prevention strategy for RVFV [36]. Adequate understanding of disease transmission is crucial to informing early warning systems [36]. For example, standard RVF outbreak risk assessments often utilise multi-criteria decision analysis tools, typically informed by expert elicitation, workshops with key decision-makers and technical experts and local climate forecasts [37, 38]. Infrequent surveillance may thus undermine the effectiveness of these early warning systems. Conversely, if uncertainty in model-derived measures is clearly communicated during the development stage, then recommendations can still be accurate, albeit potentially leading to less effective early warning systems due to the wider range of epidemiological outcomes. Our results underscore how the frequency of serological data can significantly influence model-based inference and consequently, the effectiveness of derived understanding and assessments of disease transmission.

These results were reflected in predictions of the timing and frequency of past outbreaks. In particular, cross-sectional serological surveys conducted at the end of each epidemiological year allowed for the correct identification of which years outbreaks occurred in most cases. Increased surveillance frequency beyond this refined understanding of when outbreaks happened within the year, whereas recovering basic qualitative aspects of outbreak dynamics—whether an outbreak happened or not—diminished as surveillance frequency decreased beyond annually. Annual surveillance may be sufficient for diseases which have clear seasonal drivers through a combination of environmental, entomological or sociological drivers, such as Rift Valley fever [39, 40] or bluetongue [41, 42]. However, in livestock diseases or spatial contexts which do not have these distinct annual signatures, recovering the true timing and frequency of past outbreaks may require even more frequent serological surveillance, which may present greater challenges in resource-limited settings.

The increased uncertainty in reconstructing past outbreak timing and frequency with less frequent surveillance propagated through to forecasts of the timing and size of outbreaks. Specifically, surveillance more frequent than once a year generally yielded accurate predictions of outbreak size over the next year. However, beyond this initial forecasting year, maintaining accuracy even in predicting whether an outbreak would occur required increasingly more frequent surveillance. This might not pose a significant problem in resource-rich settings where sufficiently frequent surveillance would likely negate the need for long-term forecasts, as disease monitoring would be ongoing. However, in resource-limited settings where competing priorities between diseases and non-standardised reporting procedures may lead to intermittent or irregular surveillance [43, 44], as is the case in the Comoros archipelago, the reliability of long-term predictions may be more crucial. This then raises the question of whether concentrating surveillance efforts into temporal clusters (as in the empirical data presented here) or regions with established higher levels of disease transmission offers greater benefits compared to regular, spatio-temporally dispersed surveillance, as explored in our study. Irrespective of surveillance regularity, the accuracy with which future disease transmission is predicted may thus impact assessments of disease control recommendations.

For this purpose, we also analysed how surveillance frequency impacted the effectiveness of disease control measures. This investigation revealed that the frequency of surveillance has the capacity to influence assessments of the effectiveness of disease control measures. Consequently, this impacted the recommendation of which control measure was deemed best and the level of confidence associated with that recommendation. This result aligns with previous findings demonstrating how a greater understanding of disease transmission (achieved through increased data as an outbreak progresses or more detailed livestock trade information) can enhance the accuracy of selecting appropriate livestock disease control strategies [32, 33]. This finding therefore has critical implications for control policy. Without the ability to confidently anticipate outbreaks, the effectiveness of proactive interventions, such as targeted vaccination programs, become inherently more unpredictable. This may lead to mistimed control measures, resulting in inefficient use of resources, where control measures are deployed either prematurely or too late to effectively mitigate an outbreak. It is thus important to communicate the uncertainty associated with control recommendations to policymakers, so they can reasonably weigh the benefits and risks associated with these measures.

Our study demonstrates that model-based policy recommendations are tied to the frequency of the surveillance data underpinning them, which naturally leads to questions over the wider surveillance landscape. This is highlighted by our finding that robust estimation of key epidemiological features was contingent on observing a rise in seroprevalence over the surveillance period. Consequently, the insights obtainable from cross-sectional seroprevalence surveys alone may be limited in regions with negligible disease circulation. This then motivates the need to explore what value different data types beyond the frequency of cross-sectional surveys may offer. For example, while cross-sectional (RVFV-specific IgG) seroprevalence surveys are often the standard assessment of RVFV exposure [20], longitudinal surveys have also been conducted, for example in Mayotte [45, 46], South Africa [47, 48] and Kenya [49, 50]. Longitudinal studies may encode greater information, which motivates broader questions: what is the optimal *type* of surveillance information to collect if resources allow for only one, and is there greater value in investing into multiple simultaneous surveillance efforts over a focused effort on a single type? For the latter, it may be possible to collect multiple types of other information (e.g. demographic data) alongside epidemiological surveillance; however, the benefits these additional data may offer are not always clear. For instance, age information was collected during seroprevalence surveys in the Comoros archipelago, however, obtaining accurate age data in the field can be challenging, particularly where livestock identification systems are lacking. Ultimately, answers to these questions would provide a strong rationale for improved data collection, which can enhance outbreak prediction, and in turn facilitate the planning of vaccine trials and the establishment of early warning triggers [51].

Several limitations warrant careful consideration when interpreting our epidemiological and policy insights. Our conclusions are based on five outbreak scenarios motivated by empirical data. Thus, we do not present specific control policy recommendations, but instead use these outbreak scenarios to illustrate the inferential effects of surveillance frequency; we have previously discussed control policy measures in the Comoros archipelago [15, 28]. Secondly, these outbreak scenarios are dependent on a single epidemiological model where the functional form of transmission rates is perfectly known. Empirical outbreaks will be influenced by several disease transmission mechanisms to varying degrees across different spatial contexts. The extent to which these findings are robust across different model structures is thus unknown. However, this may motivate ensemble modelling analyses for RVF and other livestock diseases, as has been achieved in human diseases [52], which can negate the impacts of specific modelling choices. One choice was to base control policy effectiveness solely on epidemiological outcomes, neglecting any economic dimension of each control measure. For policymakers, the cost-effectiveness of a strategy is a significant factor, yet quantifying this for livestock diseases is a great challenge. Furthermore, while our ranking identified optimal policy, it does not fully encapsulate the magnitude of the difference in outcome between interventions. That is, the best policy might only offer a marginal improvement over others, which may be an important aspect to consider in practical decision-making. Finally, while this study focused on the impact of surveillance frequency on predictions, it did not explicitly address the interplay with other sources of uncertainty, such as climate variability, which can also significantly impact outbreak predictability [36].

In conclusion, our study illustrates the impacts of surveillance frequency on the reliability of model-based inferences for Rift Valley fever, where less frequent surveillance increased uncertainty in epidemiological estimates, hindered accurate reconstruction of past outbreaks, and diminished the robustness of forecasts. This uncertainty directly translated into challenges in assessing the effectiveness of different disease interventions, reducing the levels of confidence in optimal control policy recommendations. Overall, our work underscores the need for optimised and sufficiently frequent surveillance to support epidemiological modelling that informs preparedness and disease control measures for livestock diseases within resource-limited settings.

## Methods

### Mathematical model

To investigate the impact of disease surveillance on model-based decision making, we first developed a stochastic, discrete-time, RVFV livestock infection model with age and spatial structure. The model was adapted from a deterministic, spatially-explicit, age-structured model [15], and is described below.

### Livestock states

The livestock population at each time *t* is described using a metapopulation framework of *K* patches, which will herein be referred to as islands. Livestock are able to move between the islands, and livestock within each island are subdivided into *A* age groups. Within each age group, livestock are further categorised by their infection status: susceptible to infection with RVFV (*S*), infected but not yet infectious (ℰ), infectious (ℐ) or recovered (ℛ). The state of a single animal at each time is thus described by their island, age group and infection status, yielding a total of 4*AK* possible states.

### Transition Matrix

Livestock transfer between states as governed by a 4*AK* × 4*AK* transition matrix *P*_*t*_, where (*P*_*t*_)_*ij*_ represents the probability of an individual transitioning from state *i* to state *j* from time *t* to time *t* + 1. The transition probability is composed of the probability of four events: survival (*f*_surv_), ageing (*f*_age_), movement (*f*_move_) and infection (*f*_inf_). Each event type depends on different components of the states *i* and *j* (the island, age group and infection status) and the current time *t*. The probability of transition is thus defined as

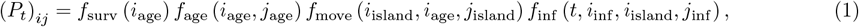

where *i*_island_, *i*_age_ and *i*_inf_ denote the island, age and infection status of any state *i*. The probability of each event type is detailed below.

It is assumed that animals in age group *a* die with an age-dependent probability denoted by *µ*_*a*_. Therefore, 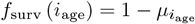. Also, animals in age group *a* aged with probability, *δ*_*a*_, so

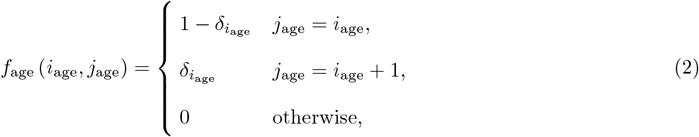

with *δ*_*A*_:= 0.

At each time step, animals move between islands in the metapopulation according to an *K × K* probability matrix *M*, where *m*_*kl*_ denotes the probability of animals moving from island *k* to island *l*. It is further assumed that only animals up to age group *A*_move_ move between islands. Therefore,

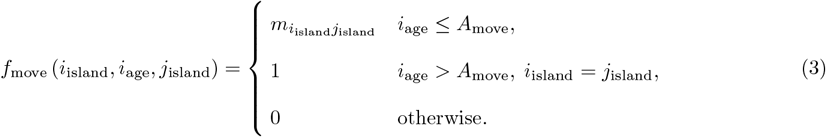

At each time *t*, susceptible animals (*S*) become infected with island-dependent probability *λ*_*tk*_. Exposed animals (ℰ) then become infectious (ℐ) with probability *ϵ* and subsequently recover (ℛ) from the disease with probability *γ* per time step. Once recovered, animals are assumed to be immune to infection for the duration of their life. The probability of transitioning from infection state *i*_inf_ to *j*_inf_ in island *i*_island_ at time *t* is thus defined as:

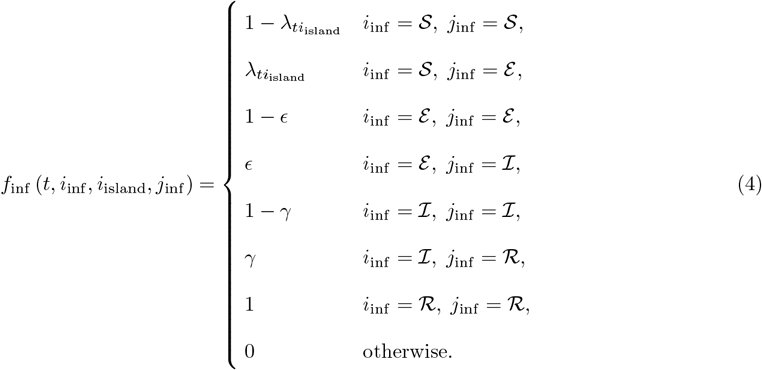

The probability of infection, *λ*_*tk*_, is assumed to linearly scale with the number of the livestock population that are infectious at time *t* in island *k* (denoted by *I*_*tk*_), and a island-dependent background level of infection *ι*_*k*_—from disease imports, wildlife reservoirs of infection and circulation within the mosquito population, for example—such that

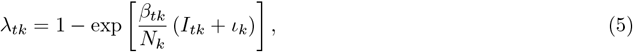

where *β*_*tk*_ denotes the disease transmission rate at time *t* in island *k* and *N*_*k*_ denotes the livestock population size on island *k*. As in Tennant et al. [15], this transmission rate scales exponentially by *α* with island-specific Normalised Difference Vegetation Index (NDVI), acting as a proxy for the effects of seasonal mosquito population dynamics on RVF disease transmission rates. Therefore,

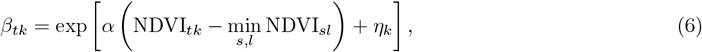

where NDVI_*tk*_ denotes the Normalised Difference Vegetation Index at time *t* on island *k*, and *η*_*k*_ denotes the (natural logarithm of the) minimum disease transmission rate in island *k*.

### Update equations

The system at time *t* is represented by a vector *X*_*t*_, where (*X*_*t*_)_*i*_ denotes the number of livestock in state *i* at time *t*. At each time step, the number of livestock in state *i* that transition to each of the possible 4*AK* states is sampled from a multinomial distribution with (*X*_*t*_)_*i*_ trials and 4*AK* + 1 events. The first 4*AK* event probabilities are given by the *i*-th row of *P*_*t*_, denoted (*P*_*t*_)_*i\**_. The final event is mortality, which occurs with probability 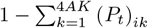. Livestock removed from the system are not tracked. Therefore, the intermediate number of individuals in each model state is given by

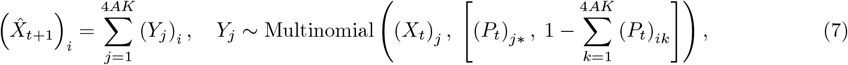

where *Y*_*j*_ represents the number of transitions from state *j* to each of the 4*AK* + 1 possible events. These intermediate model states are lastly updated such that the livestock population size of each island *k*, denoted by *N*_*k*_, is constant over time. The number of livestock in state *i* at time *t* + 1, (*X*_*t*+1_)_*i*_, is thus calculated as follows.

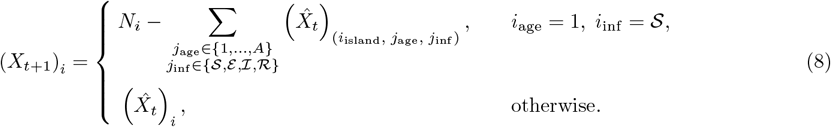

### Initialisation

At the first time point (*t* = 0), a proportion *ν*_*k*_ of the total population on each island *k* is assumed have infection status ℛ, and the remainder with infection status *S*. It is assumed that the first time point is aligned to the start of the epidemiological year, so it is assumed that there are no exposed or infectious individuals at this time point. The initial number of individuals immune to infection on island *k*, denoted *c*_*k*_, is sampled from a binomial distribution given the total population size of the island and *ν*_*k*_. That is, *c*_*k*_ ∼ Binomial (*N*_*k*_, *ν*_*k*_). It is also assumed that the initial age distribution of recovered animals was proportional to the age distribution of the total population. Therefore,

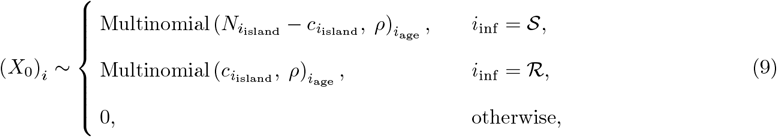

where *ρ* denotes a vector with *ρ*_*a*_ equal to the expected proportion of the livestock population in age-group *a*.

### Fixed model parameters

The parameters of the model were selected to reflect knowledge on the demography and epidemiology of RVF in the Comoros archipelago. The parameters that were fixed in the model are shown in Table 2.

**Table 2.** Fixed model parameters and priors. The fixed parameters in the metapopulation model were chosen based on the current understanding of the epidemiology of Rift Valley fever and the demography of the livestock population in the Comoros archipelago. The remaining parameters were estimated by fitting the model in a Bayesian framework to series of serological data collected across the archipelago. The fixed values and prior distribution of the model parameters are shown in the table below.

| Parameter | Description | Value / Prior |
| --- | --- | --- |
| $K$ | Number of islands in the metapopulation | 4 |
| $A$ | Number of age groups | 10 |
| $A_{\text{move}}$ | Number of age groups which can move between islands | 2 |
| $N_1$ | Total population on Grande Comore | 224 353 |
| $N_2$ | Total population on Mohéli | 31 872 |
| $N_3$ | Total population on Anjouan | 93 616 |
| $N_4$ | Total population on Mayotte | 20 052 |
| $m_{12}$ | Movement probability from Grande Comore to Mohéli | $6.5 \times 10^{-5}$ |
| $m_{13}$ | Movement probability from Grande Comore to Anjouan | $9.3 \times 10^{-6}$ |
| $m_{21}$ | Movement probability from Mohéli to Grande Comore | $8.8 \times 10^{-4}$ |
| $m_{23}$ | Movement probability from Mohéli to Anjouan | $6.5 \times 10^{-5}$ |
| $m_{31}$ | Movement probability from Anjouan to Grande Comore | $2.8 \times 10^{-4}$ |
| $m_{32}$ | Movement probability Anjouan to Mohéli | $6.7 \times 10^{-4}$ |
| $m_{34}$ | Movement probability Anjouan to Mayotte | $1.8 \times 10^{-4}$ |
| $\delta_a$ | Weekly ageing rate for age groups $a \in [1, 9]$ | 1/48 |
| $\mu_a$ | Mortality rate of animal in age groups $a \in [1, 9]$ | $8.8 \times 10^{-3}$ |
| $\mu_{10}$ | Mortality rate of animal in age group 10 | $6.2 \times 10^{-3}$ |
| $\epsilon$ | Probability of latent period ending per time step | 1 |
| $\gamma$ | Probability of recovering from infection per time step | 1 |
| $\nu_k$ | Initial proportion immune in island $k$ | Beta (5, 5) |
| $\iota_k$ | Background level of infection in island $k$ | TruncNorm (0, 0.261, 0, ∞) |
| $\eta_k$ | Minimum transmission rate in island $k$ | Normal (-2, 2) |
| $\alpha$ | Scalar for NDVI effects on disease transmission rates | TruncNorm (3, 2, 0, ∞) |
| $\phi_k$ | Observed overdispersion in island $k$ | TruncNorm (0, 1.253, 0, ∞) |

The livestock population of each island in the archipelago was described using the model, so the number of islands in the metapopulation was set to equal four (*n* = 4). On each island, ten livestock age groups were described (*A* = 10), corresponding to animals from 0–1 years old (age group 1) up to 9 years old and above (age group 10). Due to the large size of adult livestock relative to the size of the boats used to move animals between islands, only animals in the first two age groups could be moved (*A*_move_ = 2). The livestock population size of each island was calculated by aggregating cattle, sheep and goat population estimates using the Gridded Livestock Map of the World [53].

We assumed that animals could move along the network motivated by Roger et al. [54]. That is, only the following movements were permitted: Grande Comore to Mohéli, Grande Comore to Anjouan, Mohéli to Grande Comore, Mohéli to Anjouan, Anjouan to Grande Comore, Anjouan to Mohéli, and Anjouan to Mayotte. The probability of livestock moving along each path in the network was chosen based on previous consultations with the Comorian veterinary services [15] and previously reported inter-island trade estimates [24, 54]. It was further assumed that any livestock which did not move between islands remained in their current island between time steps.

The time-step for the model was an epidemiological week, which is equivalent to approximately 1.08 calendar weeks. This was done to allow simulations from the model to be easily temporally aligned with empirical seroprevalence surveys. Therefore, the weekly probability of ageing, *δ*, was set to 1*/*48 for age groups 1–9. The age distribution of livestock, *ρ*, was calculated directly from empirical data [55, 56], and corresponding age-dependent mortality rates were estimated assuming a stable age distribution over time in the absence of movement and disease [24]. The probability at which exposed livestock became infectious, *ϵ*, and the probability that infectious livestock recovered from infection, *γ*, were equal to 1, corresponding to latent and infectious periods of one week. This choice defined the seasonal reproduction number—the expected number of secondary infections caused by a single primary infection in a fully susceptible population at time *t* on island *k*—as being equal to the disease transmission rate, *β*_*tk*_ (Equation 6).

The NDVI estimates required to calculate these disease transmission rates were estimated by aggregating 16-day NDVI readings of 250m by 250m grid squares for each island [57]. These 16-day estimates were then smoothed over time using a Gaussian kernel with a standard deviation of 21 days and then extracted for each epidemiological week of the model. The remaining model parameters were inferred from serological data as described below.

### Parameter estimation

To estimate the remaining transmission model parameters, the model was fitted to age-stratified serological data in a Bayesian framework. Below we define the likelihood (observation model) and priors of each parameter that was used to fit our model.

### Observation model

For each seroprevalence survey it was assumed assumed that farms and animals were randomly sampled for serological testing on each island where within-island seroprevalence levels were spatially heterogeneous. We thus assumed that the number of RVF-specific IgG antibody positive animals of age groups *A*_obs_ on island *k* was beta-binomially distributed given (i) the total number of animals tested, (ii) the proportion of the livestock population that were immune on the island at the time of testing and (iii) an island-specific overdispersion parameter *ϕ*_*k*_. Adding overdispersion into the observation model allowed for us to account for within-island heterogeneity in livestock immunity levels, where we set an absence of overdispersion to correspond to a binomially distributed observation model as previously done by Tennant et al. [15]. The transmission model was simulated forward in time to calculate the mean proportion of livestock that were immune in age groups *a*_obs_ on island *k* during the time of the survey *t*_obs_, which is denoted by 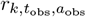. Therefore,

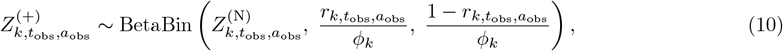

where 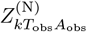 and 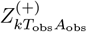 denote the number of animals that were tested and RVF antibody positive across age groups *A*_obs_ on island *k* over the time window *T*_obs_, respectively. As each survey is assumed to have been conducted independently from one another, the likelihood of jointly observing each serological survey in succession is defined by the product of observing each survey individually.

### Prior distribution

The first time point of the model corresponded to July 2008. Prior to this, (i) genomic analysis suggested the virus had been imported from East Africa during the RVF outbreak in Kenya during 2006/7 [58], (ii) human cases of RVF were detected in the archipelago in 2007/8 [59], and (iii) an elevated level of the proportion of livestock demonstrating prior infection with RVFV in Mayotte during 2007/8 was reported [45]. RVFV was thus circulating in the archipelago prior to July 2008, so moderately informative beta distribution priors with mean of 50% were set for the initial proportion of the livestock population immune on each island, *ν*. The level of background disease introduction on each island through virus maintenance in vector populations and wildlife reservoirs and imports of infected animals from outside the archipelago was assumed to be very low.

Weakly informative normal priors for *ι* were used with a mean equal to 10 background infections per year per island (in the absence of any seasonal dynamics). Weakly informative normal priors were placed on transmission parameters, *α* and *η*, as these were yet to be quantified. The levels of heterogeneity across each island were also unknown so weakly informative normal priors were used for *ϕ*. The specific prior distributions used for each parameter are stated in Table 2.

### Posterior distribution

With the definition of the likelihood and priors for parameters, we sampled from the joint posterior distribution of the parameters, *θ*, and the number of individuals in each state in the transmission model using an adaptive particle marginal Metropolis-Hastings algorithm as by Andrieu et al. [60].

In summary, this algorithm constructed a Markov chain of parameters *θ* and the number of individuals of each state in the transmission model after *T* time-steps, denoted *X*_0:*T*_. It did this by, at step *n* in the chain, first proposing a set of candidate parameters *θ*^*′*^ from a proposal distribution *q*. The candidate states 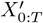 were then proposed using a particle filter. In the particle filter, a set of *S* independent simulations (particles) are initialised at *t* = 0 using the candidate parameters *θ*^*′*^. These simulations are then run forward up to the first set of seroprevalence surveys. Each simulation is given a weight according to the likelihood of observing the serological surveys under the observation model described above. Simulations are then resampled with replacement using stratified resampling [61]. This process of simulation, weighting, and resampling then progresses iteratively through the remaining serological surveys. The candidate states 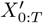 were then sampled using the final simulation weights. The mean of the final simulation weights yields an unbiased estimate of the marginal likelihood of observing data *Z*, denoted 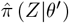 [60, 62]. The candidate pair 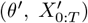 was then accepted, 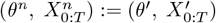, or rejected, 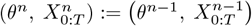, with probability

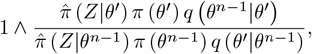

where *π* (*θ*) denotes the prior of *θ*.

The Markov chain was initialised with parameters randomly sampled from their respective prior distributions and states sampled from the particle filter process described above. All parameters were proposed jointly at each step in the chain using a multivariate normal distribution as described by Roberts and Rosenthal [63]. The number of simulations *S* to generate in the particle filter substantially affected the computational runtime and acceptance rate of candidate pairs 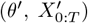. Therefore, to initially explore the parameter space and reduce computational runtime, Markov chains of 50,000 values with a low number of simulations (*S* = 20) were calculated. These chains were extended to 175,000 values whilst switching to a larger number of simulations in the particle filter (*S* = 500), increasing the acceptance rate of candidate pairs. The first 125,000 iterations of each Markov chain was then discarded as burn-in. Convergence of the chains was assessed through visual inspection of the trace plots and calculation of the revised Gelman-Rubin 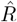 diagnostic [64].

### Surveillance frequency analysis

To assess the effect of surveillance frequency on the understanding, forecasting and control of Rift Valley fever outbreaks, we generated several series of synthetic serological surveys conducted at different time intervals under realistic outbreak scenarios inferred from empirical data. Fig 1 illustrates the process of generating serological data with different frequencies using empirical data collected across the archipelago from 2008 to 2015. Below we detail how these data were generated and subsequently used to assess the impact of a range of disease control measures.

### Inferred outbreak scenarios

Realistic outbreak scenarios were inferred by fitting the model to empirical data conducted across the archipelago from July 2009 until June 2015 (Fig 1A), where an outbreak scenario was defined as a single sample from the posterior distribution of the model fitted to the following empirical data (Fig 1B).

Cross-sectional seroprevalence surveys were conducted in 2009, 2012, 2013 and 2015 throughout the Union of Comoros—Grande Comore, Mohéli and Anjouan—as part of national surveillance programs [54, 65], and aggregated by the month in which they were conducted. Livestock seroprevalence data from the SESAM (Système d’épidémio-surveillance animale à Mayotte) surveillance system covered the period from 2008 to 2015 in Mayotte [24, 46], and were aggregated by epidemiological year, which begin in July. For surveys that were not age-specific, animals were classed as either adults (equivalent to age groups 2–10 in the model) and infants (age group 1). In surveys where the age of animals was unknown, it was assumed that these were conducted across all age groups.

### Generation and inference of synthetic serological surveys

For each of five outbreak scenarios inferred from the empirical data, synthetic serological data with different frequencies were generated (Fig 1C). At regular time intervals starting from July 2009, livestock from each island were tested for RVFV and were positive according to the observation model (Equation 10). It was assumed that each survey took place over a month testing 100 livestock—for reference, the mean sample size per survey in the empirical data was 146, 53, 71, and 539 for Grande Comore, Mohéli, Anjouan and Mayotte, respectively. It was assumed that these 100 samples were multinomially distributed across all age groups proportional to the livestock age distribution of the island at the time of the survey. Data sets with eight different surveillance frequencies were generated per outbreak scenario with one month-long surveys every 3 months, 4 months, 6 months, 1 year, 18 months, 2 years, 3 years and 6 years. To align the final dates across different surveillance frequencies, the final survey was conducted in July 2014. Across the five outbreak scenarios, this yielded a total of forty data sets.

The mathematical model was then fitted each of the forty synthetic sero-survey data sets (Fig 1D). The posterior distribution of each fitted models were compared with their respective outbreak scenario, validating the robustness of the parameter estimation procedure described above. Each fitted model was then simulated for an additional three years (up to July 2017) in the absence of disease control measures, and then the following metrics were compared across surveillance frequencies of the same outbreak scenario: (i) the posterior distribution of estimated parameters, (ii) historical and forecasted predicted weekly seroprevalence in livestock on each island and (iii) the predicted number of infections during the forecasting period.

### Evaluation of disease control measures

To illustrate the impact of surveillance frequency on model-based disease control recommendations, the model was extended to include three control measures: (i) livestock movement restrictions, (ii) disease transmission reduction and (iii) vaccination of livestock.

Livestock movement restrictions were modelled by scaling the movement matrix *M* by (1 − *κ*_move_), where *κ*_move_ represents the strength of the movement restriction. To emulate the effects of vector-control measures on reducing disease transmission, the disease transmission rate (Equation 6) was scaled by (1 − *κ*_trans_), where *κ*_trans_ represents the strength of reduction to disease transmission. Vaccination was introduced into the model by extending the state of an animal to include its vaccination status: unvaccinated or vaccinated. Unvaccinated livestock were vaccinated each year with probability *κ*_vac_. For the purposes of this study, the World Health Organization target product profile for a livestock RVFV vaccine [66] was used to set vaccine properties. That is, the probability of infection (Equation 5) for vaccinated livestock was scaled by 10%, equivalent to a vaccine efficacy of 90%, and vaccinated livestock were protected from infection in this way for a mean duration of 3 years.

Three-year forecasts were simulated from each fitted model with the following control strategies: (i) no controls, (ii) partial and complete movement restriction between islands (*κ*_move_ = {50%, 100%}), (iii) small reductions to within-island transmission rates (*κ*_trans_ = {5%, 10%, 15%}) and (iv) limited annual vaccination of livestock (*κ*_vac_ = {10%, 20%, 30%}). These strategies were then ranked in the following way.

For each surveillance frequency and outbreak scenario, we drew a single sample from the model’s joint posterior distribution. This sample was used to simulate all control measures, which were then ranked based on the predicted number of livestock infections (lower infections receiving higher ranks) over the three-year forecast. We repeated this process 2500 times, and calculated the average rank for each control strategy. Finally, the control strategies were re-ranked based on these average ranks.

### Computational implementation

The metapopulation model and fitting algorithm were coded and executed in C++20 with the GNU Scientific Library (version 2.7) [67]. The outputs of these were analysed and visualised in R (version 4.4.1) [68] and the tidyverse library (version 2.0.0) [69]. The datasets used to fit and simulate the mathematical model forward in time, a full description of the data, all code, including the metapopulation model, fitting algorithm, and data analysis scripts, and a walk through of how to replicate our analysis for one outbreak scenario and seroprevalence survey frequency in our study are fully available through the GitHub repository: wtennant/rvf surveillance. [70].

## Supporting information

**S1 Fig.**
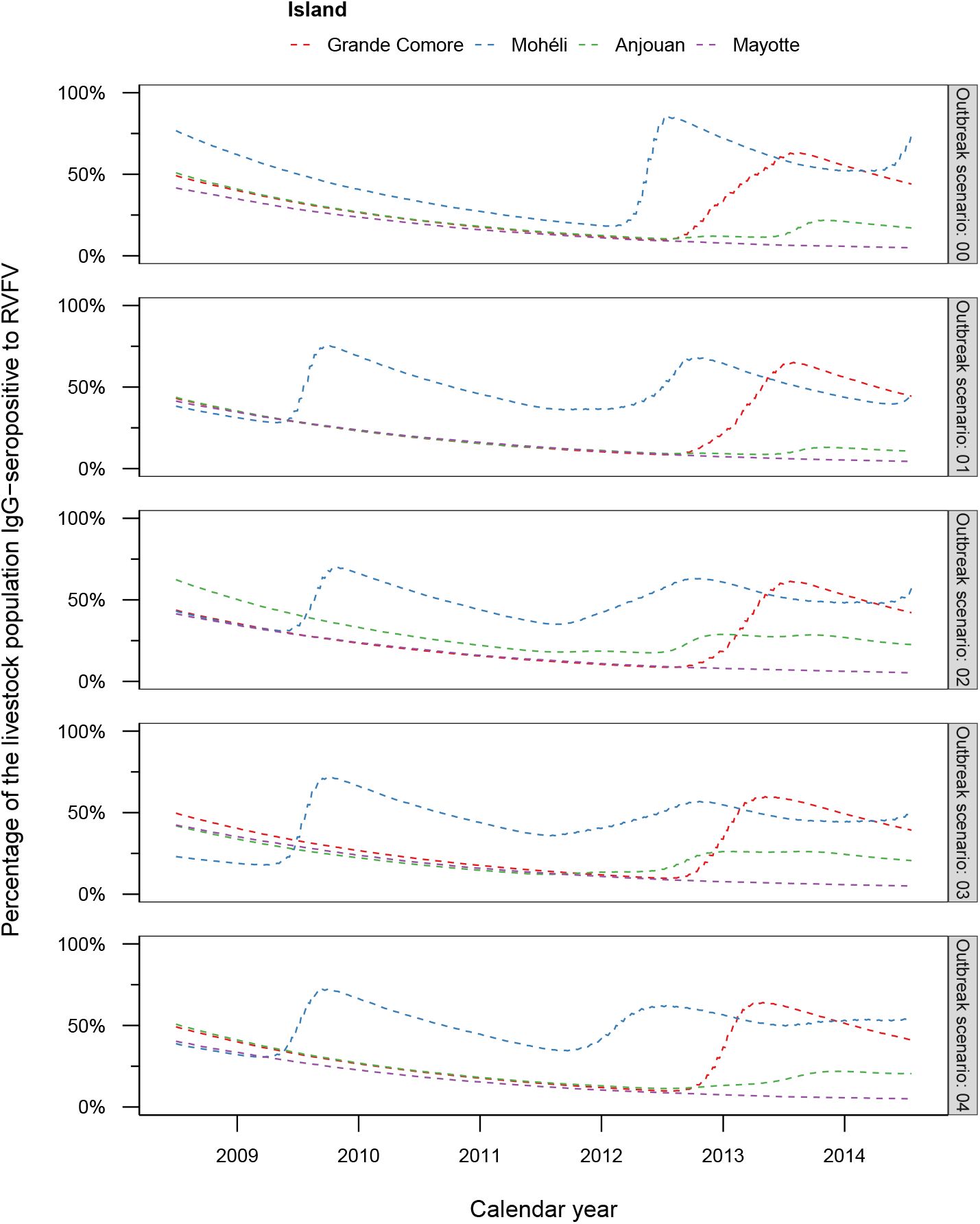
Outbreak scenarios. The mathematical model was fitted to empirical seroprevalence data conducted from July 2008 to June 2015 across the Comoros archipelago—Grande Comore (red), Mohéli (blue), Anjouan (green) and Mayotte (purple). Five outbreak scenarios were extracted from the fitted model. Shown is the modelled percentage of livestock that are RVFV-specific immunoglobulin-G (IgG) seropositive, or the modelled percentage of livestock which have prior RVFV infection, on each island in the archipelago for the five outbreak scenarios.

**S2 Fig.**
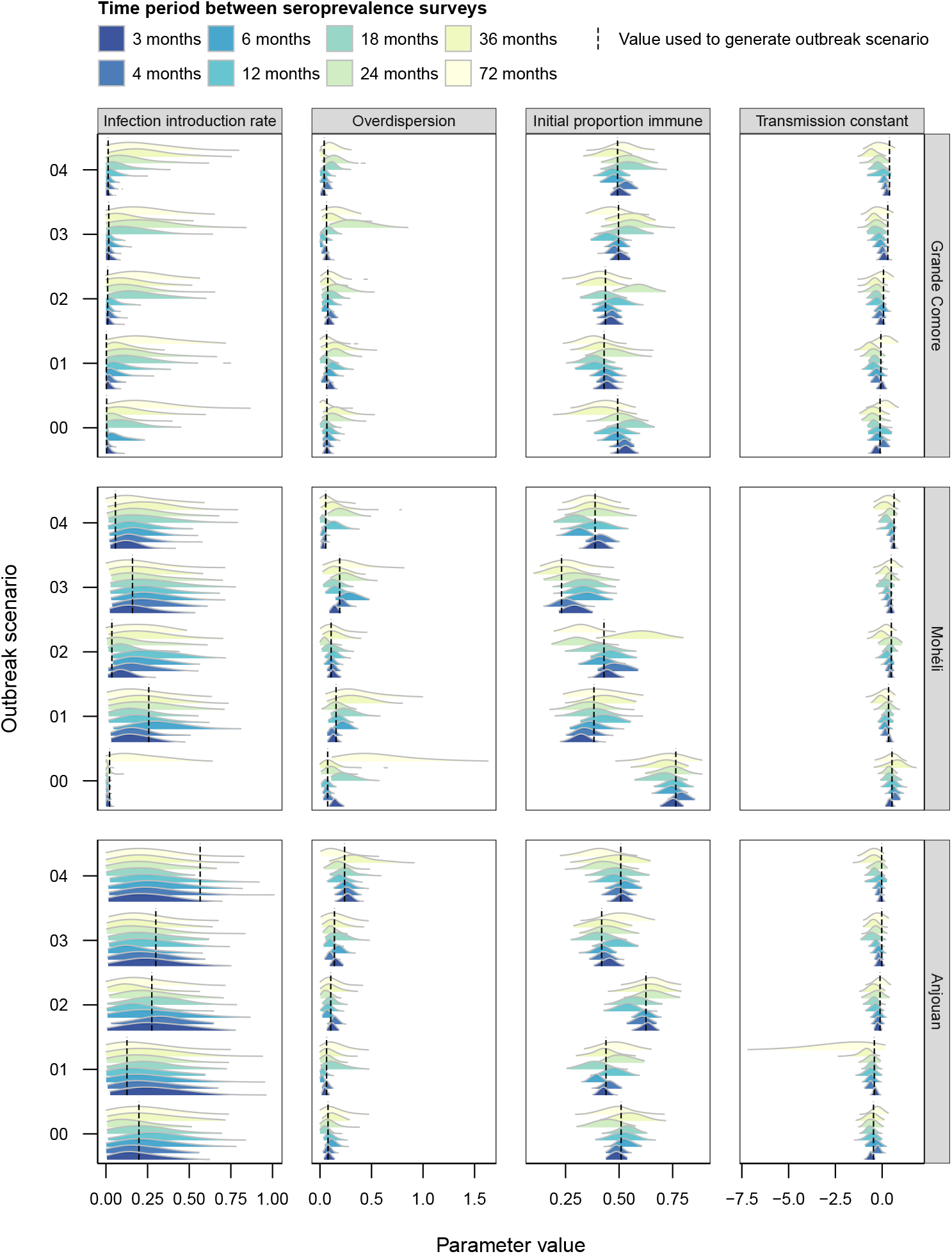
Posterior distribution of parameters for Grande Comore, Mohéli and Anjouan for each surveillance frequency and outbreak scenario. Each outbreak scenario was used to generate seroprevalence surveys conducted at different frequencies: one month-long survey every three months, four months, six months, year, two years, three years and six years (from dark blue to cream). Model parameters were then estimated for each surveillance frequency, and compared with the value used to generate them (vertical black dashed line). Shown are density estimates of each parameter for Grande Comore, Mohéli and Anjouan for each surveillance frequency and outbreak scenario. The density estimates are based on 5000 samples from the posterior distribution of each fitted model.

**S3 Fig.**
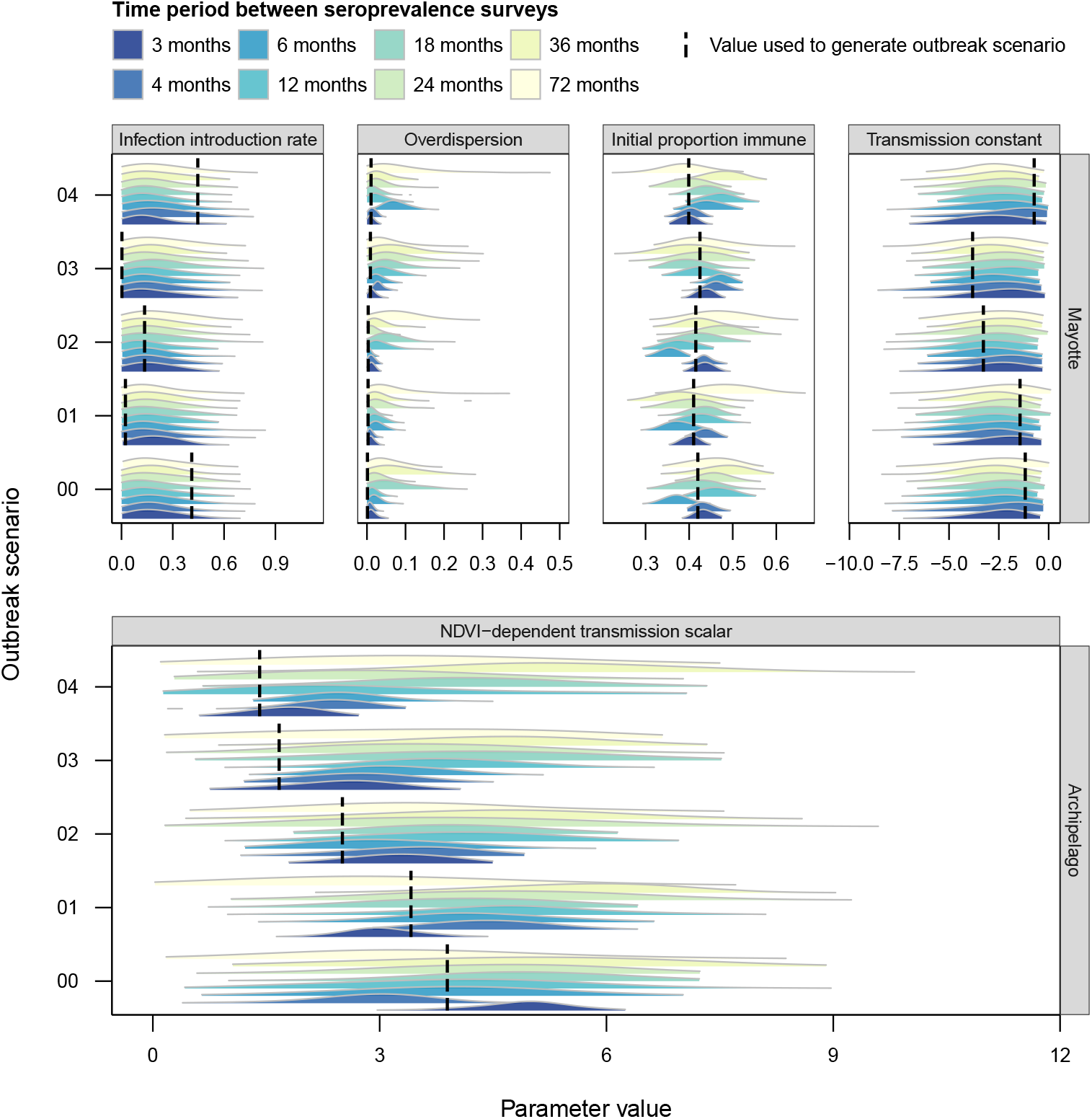
Posterior distribution of parameters for Mayotte and parameters shared across the archipelago for each surveillance frequency and outbreak scenario. Each outbreak scenario was used to generate seroprevalence surveys conducted at different frequencies: one month-long survey every three months, four months, six months, year, two years, three years and six years (from dark blue to cream). Model parameters were then estimated for each surveillance frequency, and compared with the value used to generate them (vertical black dashed line). Shown are density estimates of each parameter for Mayotte, and for the parameter shared across all islands in the archipelago. The density estimates are based on 5000 samples from the posterior distribution of each fitted model.

**S4 Fig.**
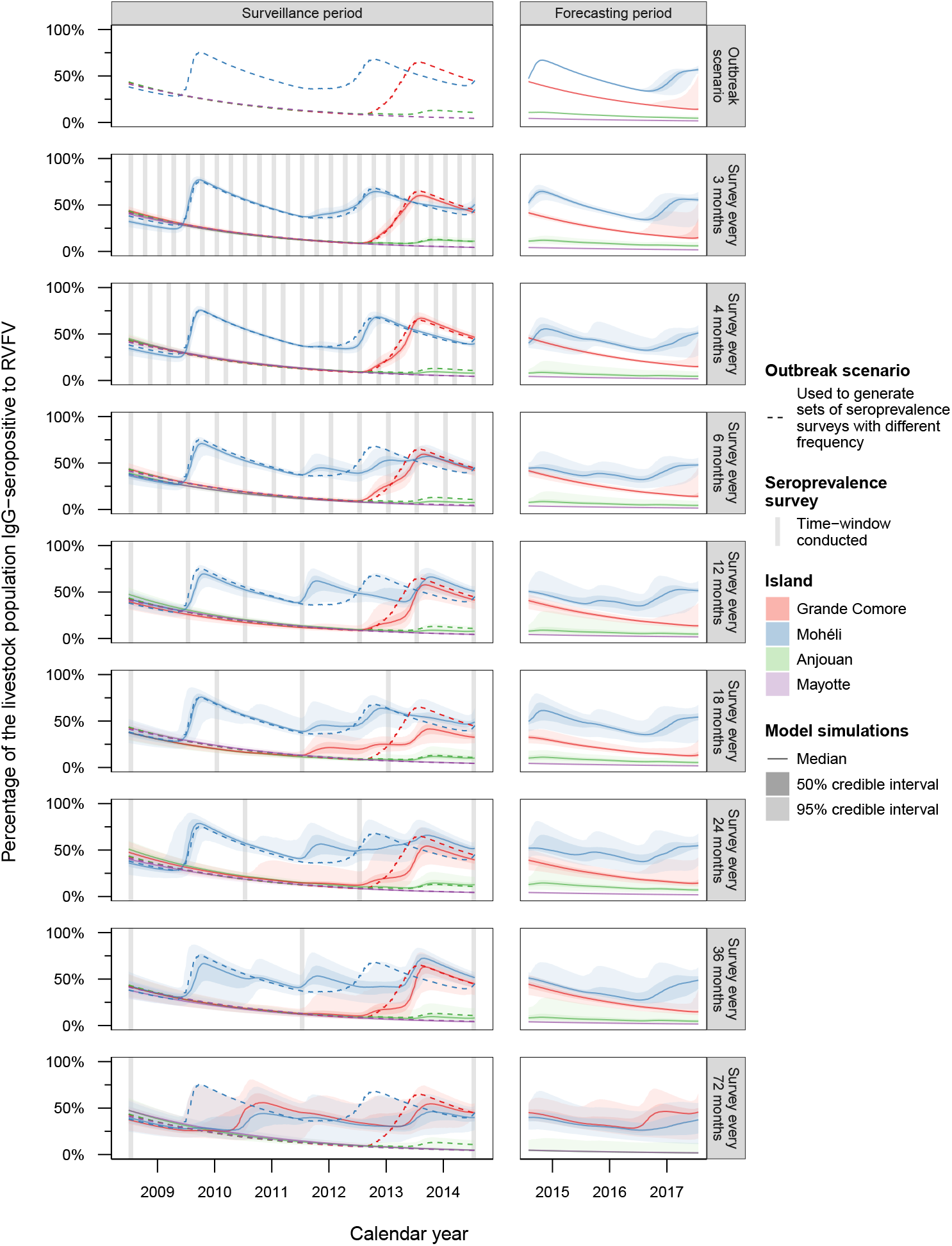
Historical and forecasted IgG-seroprevalence to RVFV (outbreak scenario 01). An outbreak scenario (dashed line) was generated by fitting a mathematical model to empirical age-stratified seroprevalence surveys conducted across the archipelago from July 2008 to June 2015. From this, synthetic serological surveys were generated with different intervals (vertical grey bars): from once every 3 months to once every 6 years. The model was fitted independently to each of these data. Shown is the median (solid line), 50% credible interval and 95% credible interval (shaded ribbons) of the percentage of livestock with past exposure to RVFV on each island in the Comoros arhcipelago—Grande Comore (red), Mohéli (blue), Anjouan (green) and Mayotte (purple). The medians and credible intervals were based on 2500 posterior samples and corresponding forward-simulations for each surveillance frequency.

**S5 Fig.**
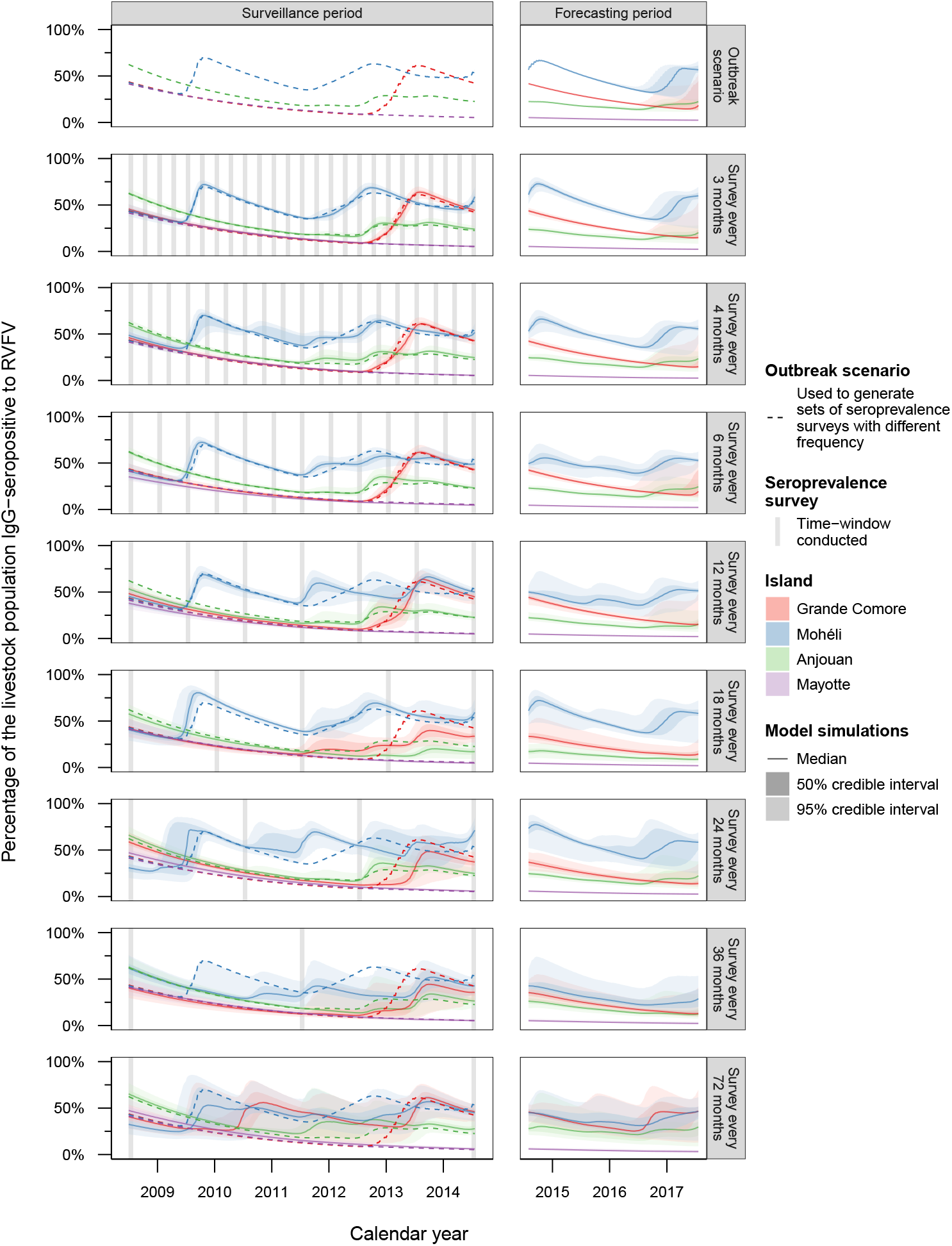
Historical and forecasted IgG-seroprevalence to RVFV (outbreak scenario 02). An outbreak scenario (dashed line) was generated by fitting a mathematical model to empirical age-stratified seroprevalence surveys conducted across the archipelago from July 2008 to June 2015. From this, synthetic serological surveys were generated with different intervals (vertical grey bars): from once every 3 months to once every 6 years. The model was fitted independently to each of these data. Shown is the median (solid line), 50% credible interval and 95% credible interval (shaded ribbons) of the percentage of livestock with past exposure to RVFV on each island in the Comoros arhcipelago—Grande Comore (red), Mohéli (blue), Anjouan (green) and Mayotte (purple). The medians and credible intervals were based on 2500 posterior samples and corresponding forward-simulations for each surveillance frequency.

**S6 Fig.**
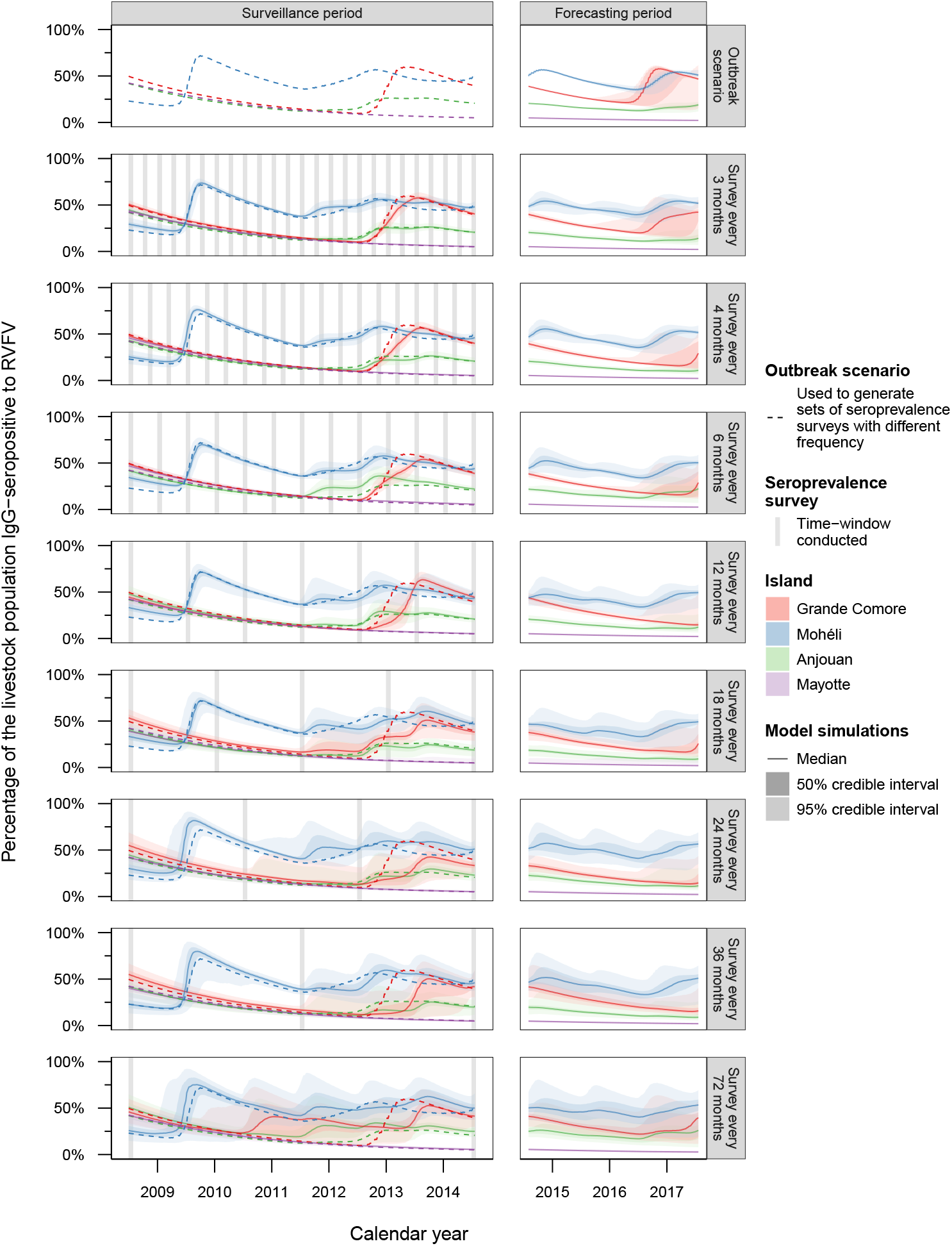
Historical and forecasted IgG-seroprevalence to RVFV (outbreak scenario 03). An outbreak scenario (dashed line) was generated by fitting a mathematical model to empirical age-stratified seroprevalence surveys conducted across the archipelago from July 2008 to June 2015. From this, synthetic serological surveys were generated with different intervals (vertical grey bars): from once every 3 months to once every 6 years. The model was fitted independently to each of these data. Shown is the median (solid line), 50% credible interval and 95% credible interval (shaded ribbons) of the percentage of livestock with past exposure to RVFV on each island in the Comoros arhcipelago—Grande Comore (red), Mohéli (blue), Anjouan (green) and Mayotte (purple). The medians and credible intervals were based on 2500 posterior samples and corresponding forward-simulations for each surveillance frequency.

**S7 Fig.**
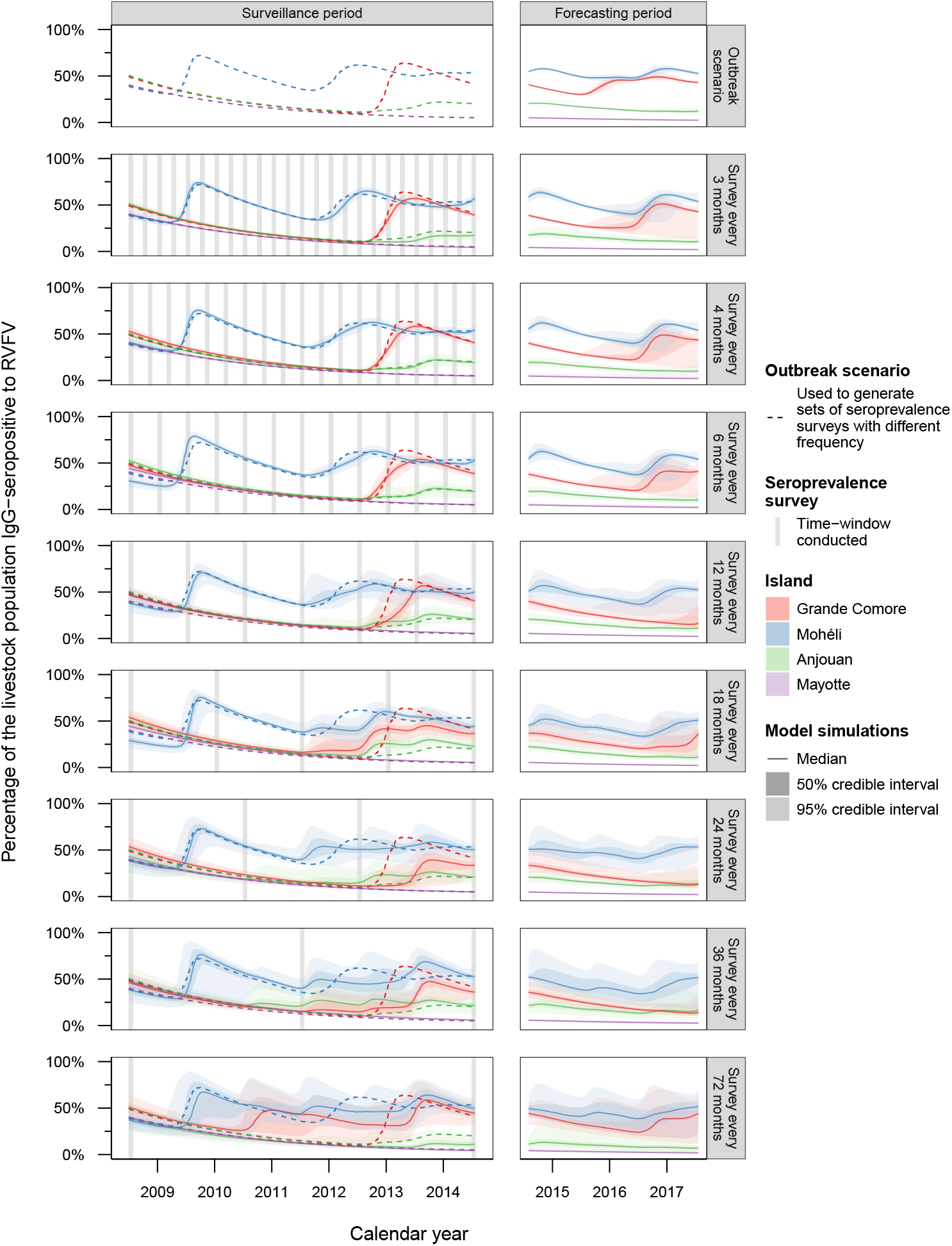
Historical and forecasted IgG-seroprevalence to RVFV (outbreak scenario 04). An outbreak scenario (dashed line) was generated by fitting a mathematical model to empirical age-stratified seroprevalence surveys conducted across the archipelago from July 2008 to June 2015. From this, synthetic serological surveys were generated with different intervals (vertical grey bars): from once every 3 months to once every 6 years. The model was fitted independently to each of these data. Shown is the median (solid line), 50% credible interval and 95% credible interval (shaded ribbons) of the percentage of livestock with past exposure to RVFV on each island in the Comoros arhcipelago—Grande Comore (red), Mohéli (blue), Anjouan (green) and Mayotte (purple). The medians and credible intervals were based on 2500 posterior samples and corresponding forward-simulations for each surveillance frequency.

**S8 Fig.**
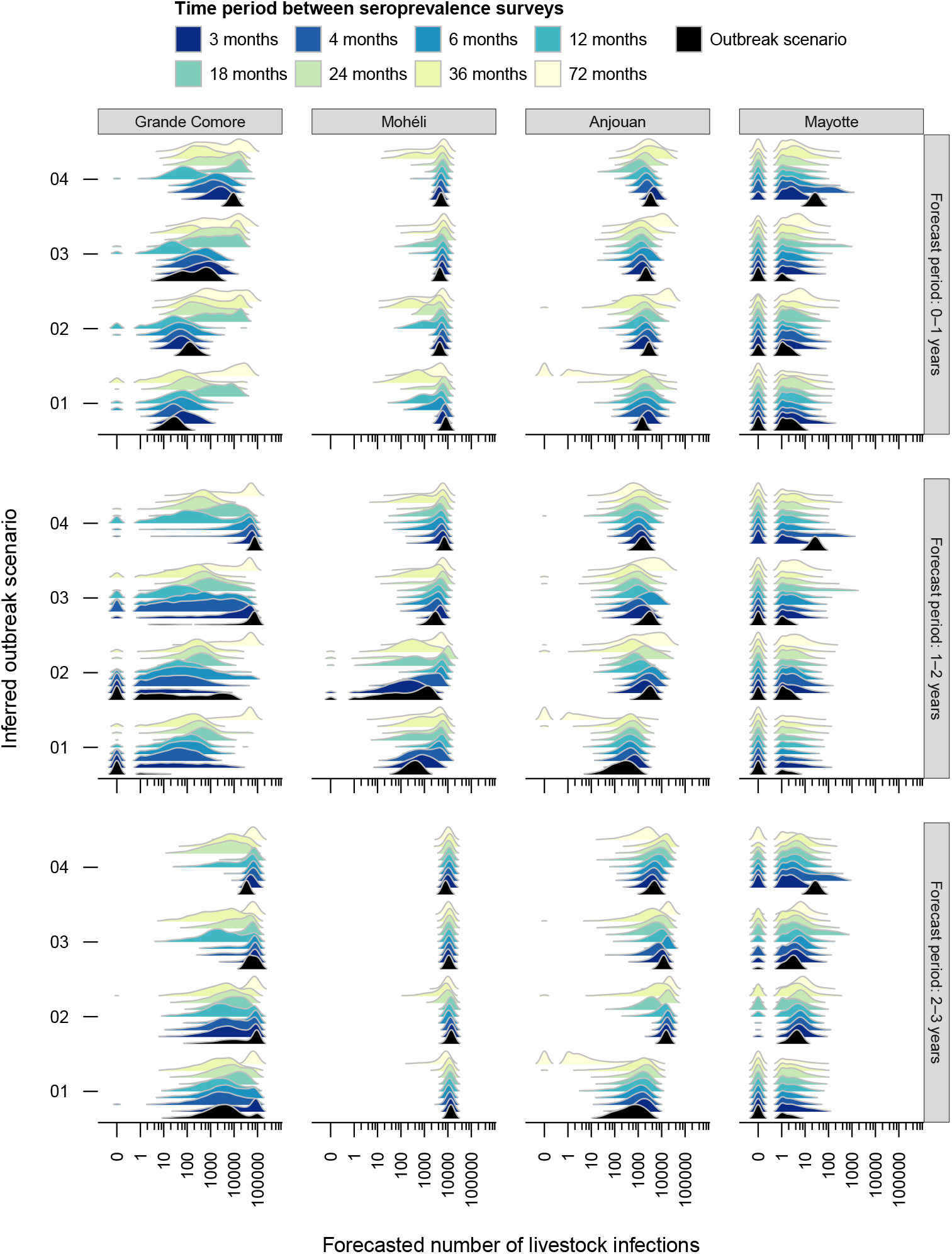
Surveillance frequency influence on outbreak size forecasts for alternate outbreak scenarios. The model was simulated forward in time for three-years without disease control measures. Shown is the predicted number of livestock infections for each year following the end of disease surveillance, which were conducted every 3 months (dark blue) to six years (light yellow), the latter four outbreak scenarios. The estimated distribution of infections from each outbreak scenario are shown in black. Estimates shown were based on 2500 simulations for each surveillance frequency from each outbreak scenario.

**S9 Fig.**
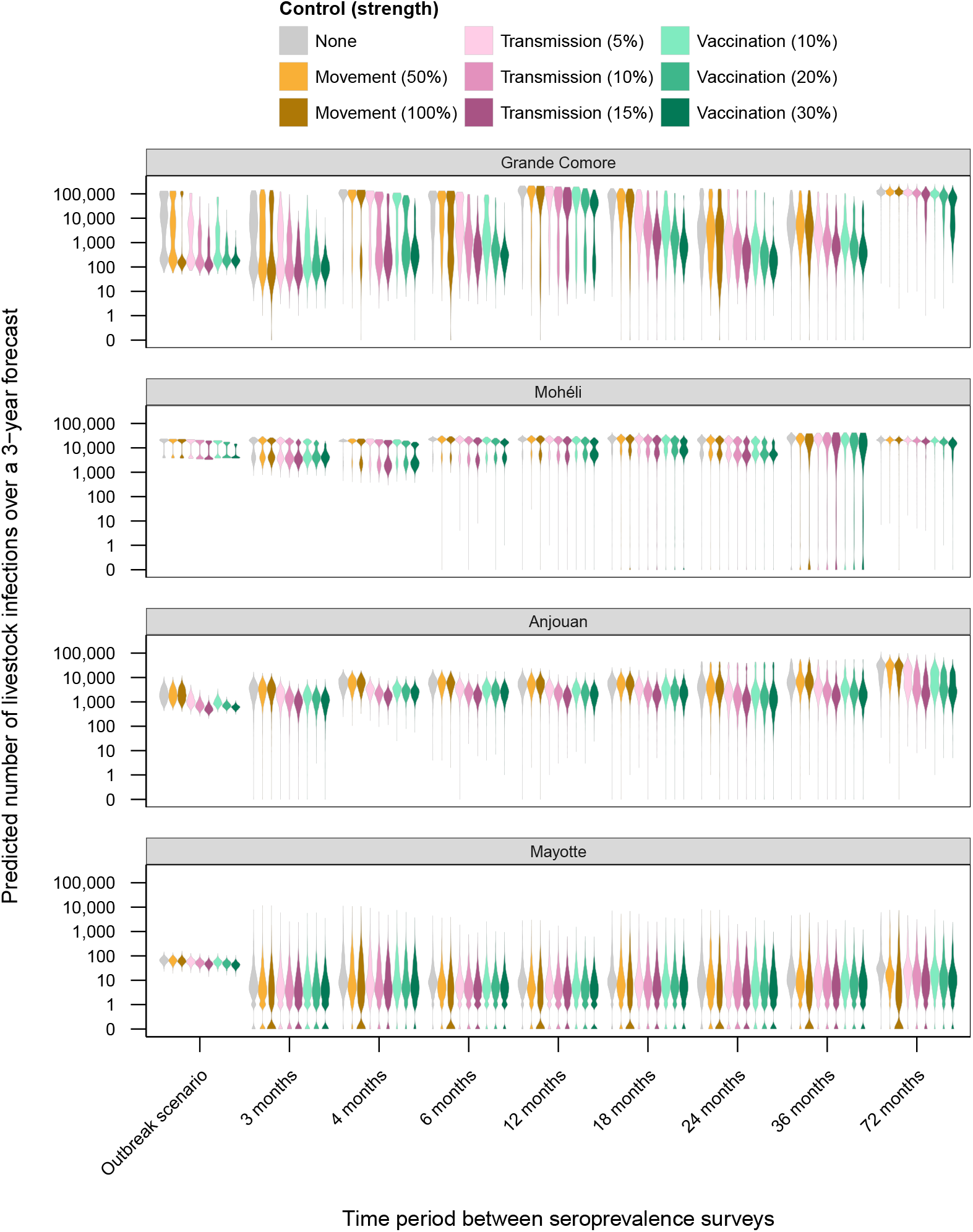
Model-predicted livestock infections under disease control measures (outbreak scenario 00). Three-year forecasts were simulated for each surveillance frequency under the following control strategies: no controls (grey), partial and complete movement restriction between islands (yellow), small reductions to within-island transmission rates (pink) and limited annual vaccination of livestock (green). Shown is the distribution of the predicted number of livestock infections over the three-year forecast for each surveillance frequency and control strategy. Distributions were based on 2500 simulations from the posterior distribution of fitted models.

**S10 Fig.**
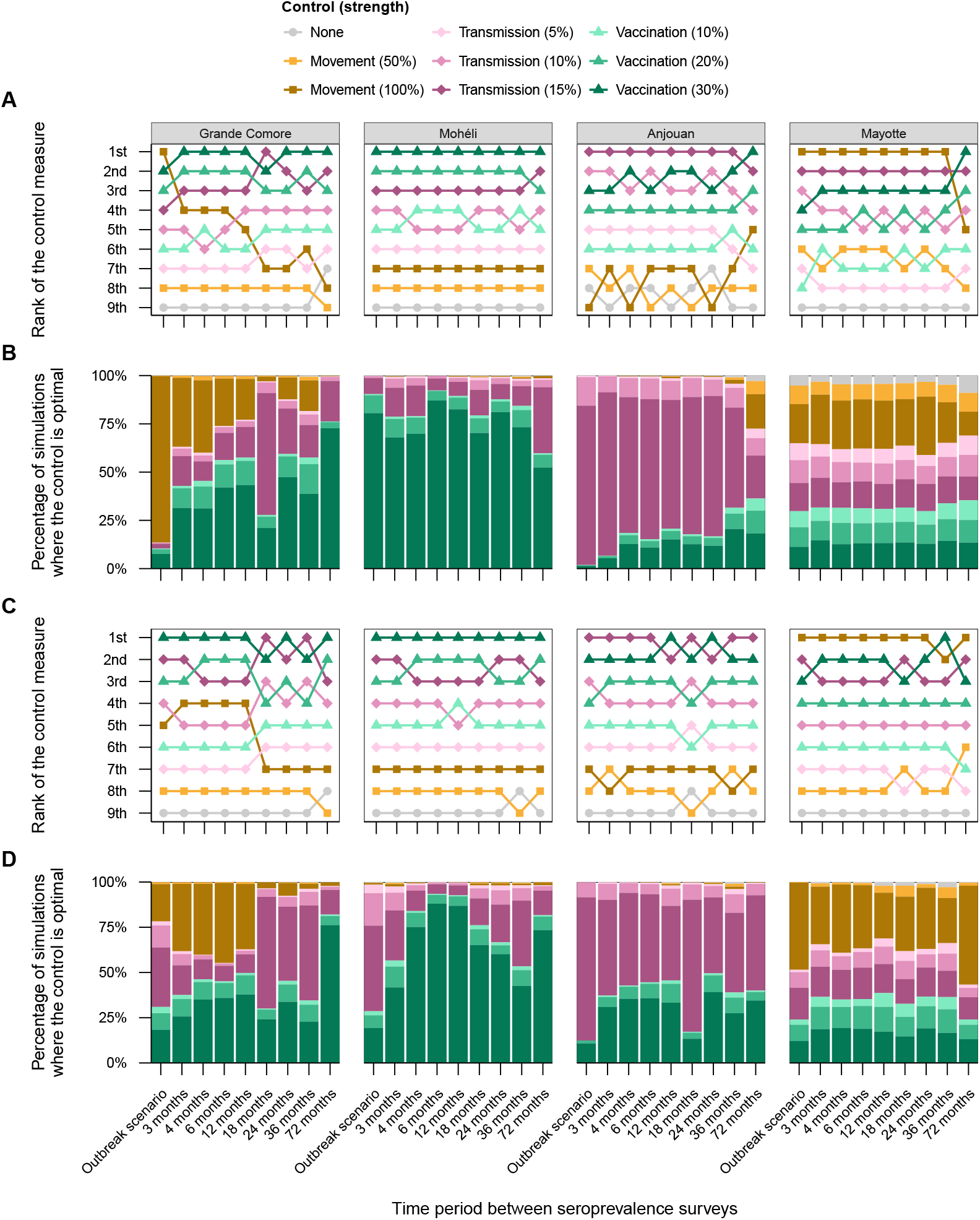
Effects of surveillance frequency on the effectiveness and confidence level in disease control strategies (outbreak scenarios 01–02). For each surveillance frequency, the number of livestock infections over a 3-year period was forecasted without (grey) and with different control measures: movement (yellow squares), reduction to transmission (pink diamonds) and vaccination (green triangles). **(A, C)** The control strategies were ranked based upon the total infections predicted within each island (lower predictions receiving higher ranks). **(B, D)** The proportion of simulations where each control strategy was the best (ranked first). Model forecasts were based on 2500 simulations for each control measure and surveillance frequency.

**S11 Fig.**
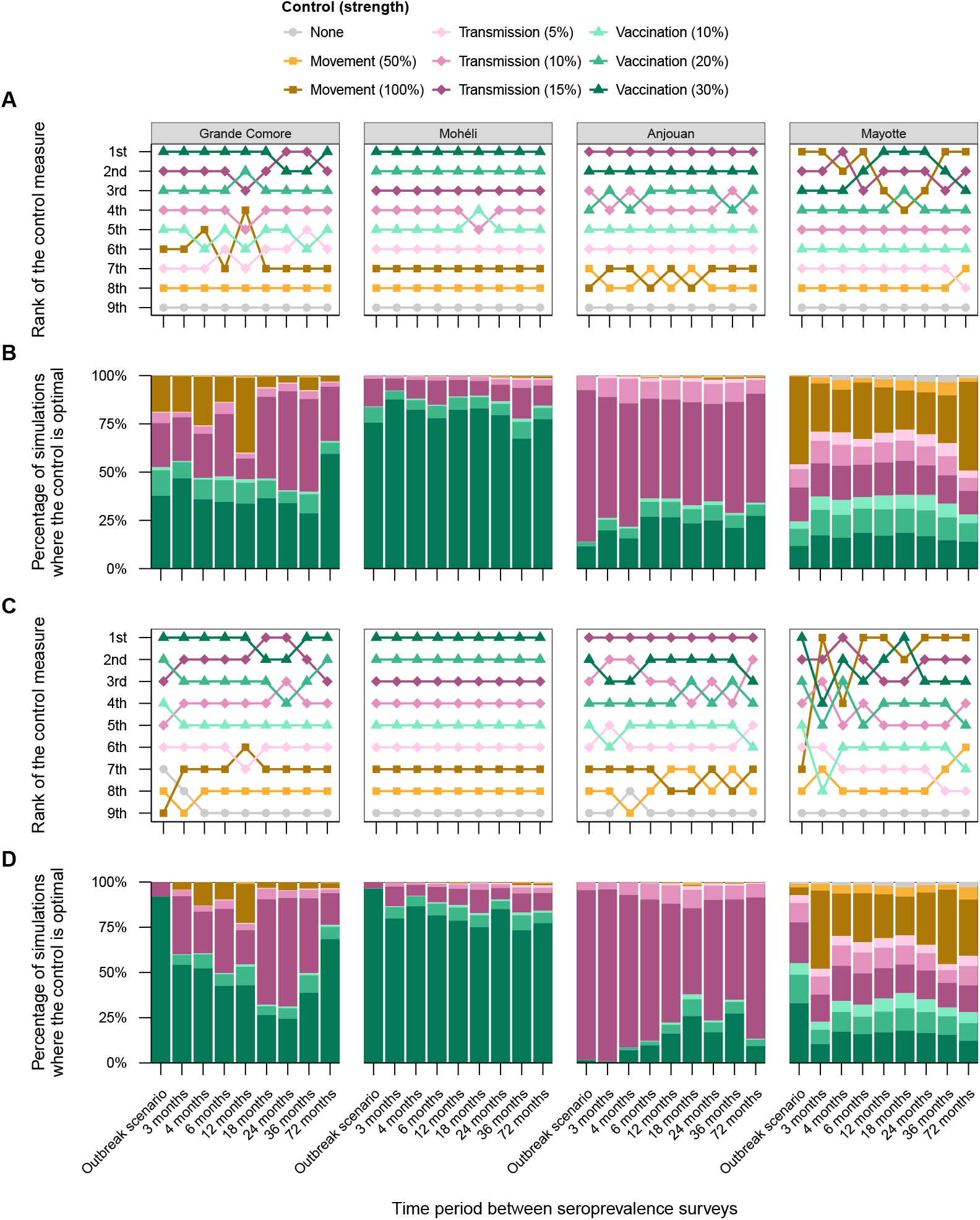
Effects of surveillance frequency on the effectiveness and confidence level in disease control strategies (outbreak scenarios 03–04). For each surveillance frequency, the number of livestock infections over a 3-year period was forecasted without (grey) and with different control measures: movement (yellow squares), reduction to transmission (pink diamonds) and vaccination (green triangles). **(A, C)** The control strategies were ranked based upon the total infections predicted within each island (lower predictions receiving higher ranks). **(B, D)** The proportion of simulations where each control strategy was the best (ranked first). Model forecasts were based on 2500 simulations for each control measure and surveillance frequency.

